# Calretinin and calbindin architecture of the midline thalamus associated with prefrontal-hippocampal circuitry

**DOI:** 10.1101/2020.07.21.214973

**Authors:** Tatiana D. Viena, Gabriela E. Rasch, Daniela Silva, Timothy A. Allen

**Affiliations:** Cognitive Neuroscience Program, Department of Psychology, Florida International University, Miami, FL, 33199, USA; Department of Environmental Health Sciences, Robert Stempel College of Public Health, Florida International University, Miami, FL, 33199, USA

**Author notes:** Corresponding Author: Timothy A. Allen, PhD, Department of Psychology, Florida International University, 11200 SW 8^th^ Street, Miami, FL, 33199, Website: http://allenlab.fiu.edu/, Twitter: @AllenNeuroLab. **Authors Contributions**, TDV and TAA designed experiments; TDV, GER, and DS performed research; TDV drafted the manuscript; TDV and GER analyzed the results; and TDV, GER, DS, and TAA edited the manuscript.

**Keywords:** calretinin, calbindin, parvalbumin, nucleus reuniens, paratenial, rhomboid, paraventricular nucleus

## Abstract

The midline thalamus bi-directionally connects the medial prefrontal cortex (mPFC) and hippocampus (HC) creating a unique cortico-thalamo-cortico circuit fundamental to memory and executive function. While the anatomical connectivity of midline thalamus has been thoroughly investigated, little is known about its cellular organization within each nucleus. Here we used immunohistological techniques to examine cellular distributions in the midline thalamus based on the calcium binding proteins parvalbumin (PV), calretinin (CR), and calbindin (CB). We also examined these calcium binding proteins in a population of reuniens cells known to project to both mPFC and HC using a dual fluorescence retrograde adenoassociated virus (AAV) based tracing approach. These dual reuniens mPFC-HC projecting cells, in particular, are thought to be important for synchronizing mPFC and HC activity. First, we confirmed the absence of PV^+^ neurons in the midline thalamus. Second, we found a common pattern of CR^+^ and CB^+^ cells throughout midline thalamus with CR^+^ cells running along the nearby third ventricle (3V) and penetrating the midline. CB^+^ cells were consistently more lateral and toward the middle of the dorsal-ventral extent of the midline thalamus. Notably, single-labeled CR^+^ and CB^+^ zones were partially overlapping and included dual-labeled CR^+^/CB^+^ cells. Within RE, we also observed a CR and CB subzone specific diversity. Interestingly, dual mPFC-HC projecting neurons in RE expressed none of the calcium binding proteins examined, but were contained in nests of CR^+^ and CB^+^ cells. Overall, the midline thalamus was well organized into CR^+^ and CB^+^ rich zones distributed throughout the region, with dual mPFC-HC projecting cells in reuniens representing a unique cell population. These results provide a cytoarchitectural organization in the midline thalamus based on calcium binding protein expression, and sets the stage for future cell-type specific interrogations of the functional role of these different cell populations in mPFC-HC interactions.

## INTRODUCTION

Interactions between the rodent agranular medial prefrontal cortex (mPFC) and the hippocampus (HC) are essential to cognition and adaptive behavior, especially the flexible use and consolidation of memory (Churchwell & Kesner, 2011; Eichenbaum, 2017; Jin & Maren, 2015; McGaugh et al., 2019; Preston & Eichenbaum, 2013). Theoretically, mPFC-HC dysfunction is a common root cause of various often overlapping neurocognitive symptoms that define several mental health disorders including Alzheimer’s disease (Braak & Braak, 1991), schizophrenia (Lisman et al., 2010), epilepsy (Gelinas et al., 2016), and others. Anatomically, mPFC-HC interactions occur through multiple circuits including direct ventral HC → mPFC projections (Cenquizca & Swanson, 2007; Ferino et al., 1987; Skelin et al., 2019; Spellman et al., 2015); indirect cortico-cortico pathways via entorhinal (Burwell, 2000; Kerr et al., 2007; Witter et al., 2017), perirhinal cortex (Furtak et al., 2007; Jayachandran et al., 2019) and retrosplenial cortex (Hunsaker & Kesner, 2018; Nelson et al., 2014); and through midline thalamo-cortical connections (Hoover & Vertes, 2012; Vertes et al., 2007, 2015). In this regard, the midline thalamus is unique in that it serves a fundamental role in creating the canonical higher-order cortico-thalamo-cortico circuitry that unites mPFC with the HC (Sherman, 2017; Dolleman-van der Weel et al., 2019).

Major subdivisions of the midline thalamus include the dorsally situated paraventricular (PVT) and parataenial (PT) nuclei, and the ventrally situated rhomboid (RH) and reuniens (RE). Each region has unique connectivity patterns with mPFC, the HC, entorhinal and perirhinal cortex and septum, and including special dual mPFC-HC projecting neurons (Su & Bentivoglio, 1990; Varela et al., 2014; Vertes et al., 2006; Vertes & Hoover, 2008). Specific anatomical connectivity patterns have been the primary contributor in understanding and predicting subregional differences (Dolleman-van der Weel & Witter, 1996; Vertes et al., 2006, 2015). For example, PVT has more connectivity with the amygdala (Li & Kirouac, 2008; Su & Bentivoglio, 1990) and been related to adaptive fear memory (Choi & McNally, 2017; Penzo et al., 2015). While RE, the most studied region, contains numerous cells with bidirectional monosynaptic projections to CA1 and mPFC (Hoover & Vertes, 2012; Varela et al., 2014), and thus been related to memory consolidation (Barker & Warburton, 2018; Dolleman-van der Weel et al., 2019; Varela et al., 2014), top-down memory functions (Ito et al., 2015; Jayachandran et al., 2019; Viena et al., 2018; Xu & Sudhof, 2013), and mPFC-HC synchrony (Ferraris et al., 2018; Hallock et al., 2016; Hauer et al., 2019; Roy et al., 2017)

Little is known about the organization of cell types in the midline thalamus. Recently, Lara-Vasquez et al. (2016) identified two populations of midline thalamic cells that were differentiated by their calcium binding protein expression, notably and calretinin (CR) and calbindin (CB). Calcium binding proteins regulate several extra- and intra-cellular functions such as cell-cell communication, cell contracture, and signal transduction by triggering or buffering calcium signaling and compartmental concentrations (Arai et al., 1994; Celio, 1990; del Río & DeFelipe, 1996; Fonseca & Soriano, 1995; Sloviter, 1989; Zimmer et al., 1995). Lara-Vazquez et al. (2016) went on to show that the *in vivo* activity of midline thalamic neurons were determined, in part, by their calcium binding protein expression. Specifically, they showed that (during urethane anesthesia) midline thalamic CR^+^ cells fired at low rates, did not increase their activity during HC theta, and were inhibited during HC sharp-wave ripples. Conversely, CR^-^ cells fired faster and responded to HC theta.

In the rat brain, the anatomical distribution of calcium binding proteins, including parvalbumin (PV), CR and CB, differ widely across brain regions. For example, in the neocortex, PV positive (PV^+^) and CB^+^ neurons are found throughout layers II-V, while CR positive (CR^+^) cells are mainly found in superficial cortical layers (Condé et al., 1994; DeFelipe, 1997; Hof et al., 1999; Reynolds et al., 2004; Sloviter, 1989). In the HC, PV^+^ neurons can be found in restricted layers and neuronal types in CA1 and CA3 such as non-pyramidal basket and axo-axonic cells, while CB^+^ cells are localized in CA1 and CA2 (Kosaka et al., 1988; Aika et al., 1994; Fonseca and Soriano, 1995; Fuchs et al., 2007), and CR^+^ neurons preferentially stain interneurons in CA1 (Miettinen et al., 1992; del Río and DeFelipe, 1996; Gulyás et al., 1996; Urbán et al., 2002). In the thalamus, calcium binding protein distributions appear highly specific across nuclei. Early studies showed that the midline thalamus labels particularly strongly for CR and CB, but not PV (Arai et al., 1994; Winsky et al., 1992). However, regional and sub-regional distributions of these calcium binding proteins in the midline thalamus have not been explored in detail.

Specifying the expression and topography of calcium binding proteins in midline thalamus will provide novel insights into structure and provide opportunities for cell-type specific targeting. Here, we used immunohistological techniques to label and analyze details of the distribution PV, CR and CB throughout the midline thalamus. We also targeted calcium binding protein expression specifically in populations of RE cells that project to both mPFC and HC using dual retrograde viral tracers, in combination with immunohistochemistry. Our results first confirmed the absence of PV^+^ cells throughout the midline thalamus. Second, we showed that there are distinct functional zones in midline thalamus defined by the topography of CR^+^ and CB^+^ labeling. Notably, we observed that the pattern of CR^+^ and CB^+^ zones were well matched between dorsal midline thalamus (PVT and PT) and ventral midline thalamus (RH and RE), only they were inverted relative to each other reflecting their wrapping around with the third ventricle (3V). CR^+^ zones were most dense medial-laterally against the dorsal or ventral 3V walls and ran dorso-ventrally, or ventro-medially, along the midline away from their respective 3V. CB^+^ cells were clustered more ventro- or dorso-laterally (for dorsal and ventral midline thalamus, respectively). CR^+^ and CB^+^ zones also overlapped and contained dual-labeled CR^+^/CB^+^ cells. Lastly, we show that dual mPFC-HC projecting cells in RE expressed none of the three calcium binding proteins examined (PV, CR, and CB) but were surrounded by dense nests of CR^+^ and CB^+^ cells. We discuss these results with respect to the mPFC-HC circuitry, and the implications of future functional interrogations of the midline thalamus.

## MATERIALS AND METHODS

### Animal care and use

All procedures described were conducted in compliance with Florida International University (FIU) Institutional Animal Care and Use Committee (IACUC) and Institutional Biosafety Committee (IBC). Brain tissue sections (40-μm thick) from a total of 16 Long Evans rats (14 males, 2 females; Charles River; 250-350g on arrival) were used in these experiments. Rats were housed individually in a 12 hours inverse light/dark cycle (lights off at 10 a.m.) and had ad libitum access to food and water.

### Retrograde tracer injections

A subsample of rats (n = 4, 2 males and 2 females) received bilateral retrograde AAV-CAG-TdTomato (59462-AAV_rg_; Addgene, MA) in mPFC (PL/IL) at the following DV coordinates: −5.0mm (50nL), −4.4mm (150 nL), and −3.8mm (200 nL), and AAV-CAG-GFP (Addgene, MA; 37825-AAV_rg_) targeting vCA1 at −7.2 mm (100 nL), 6.8 mm (200 nL), and 6.2 mm (200 nL). Retrograde viral vector expression post injections was between 6-8 weeks before animals were sacrificed. Thalamic brain tissue sections from these rats further underwent 3,3’-diaminobenzidine (DAB) reactions of CR or CB (see below) prior to image visualization and captures using an Olympus BX41 brightfield/epifluorescence microscope.

### Immunohistochemical tissue processing

Naïve and experimental rats were deeply anesthetized and transcardially perfused with 100 mL of heparin saline at a speed of 10mL/minute, followed by 250 mL of 4% paraformaldehyde (PFA, pH 7.4) at the same perfusion speed. Post perfusion, brains were removed, preserved in 4% PFA for 24 hours and then cryoprotected in 30% sucrose solution until they sank to the bottom. Subsequently, fixated brains were frozen and cut into coronal sections using a cryostat (Leica CM 3050S) or a sliding microtome.

All tissue sections were cleaned with 1% sodium-borohydride in 0.1 M PB (pH 7.4), blocked for 1 hour in 0.5% Bovine Serum Album (BSA) and then processed using the following procedures:

### Parvalbumin fluorescence reactions

A set of thalamic brain sections from 4 male rats was incubated at room temperature for 48 hours in parvalbumin (PV) primary antibody (1:250, MCA-3C9; Encore Bio, FL). After washes, tissue was incubated for 5 hours at room temperature in VectaFluor DyLight 594 anti-mouse secondary antibody (3 drops in 5mL of 0.1% BSA, DK-8818; Vector Labs, CA). When incubation was completed, tissue was washed in 0.1 M PB (3 x 5 minutes) then mounted on gelatin coated slides and coverslipped with VectaShield mounting medium with DAPI for visualization.

### Calretinin and calbindin dual fluorescence reactions

Thalamic brain tissue sections from 5 male rats were incubated for 48 hours at room temperature in mouse CB primary antibody (1:500, MCA-5A9; Encore Bio, FL) and 24 hours in rabbit CR primary antibody (1: 2000, RPCA-Calret; Encore Bio, FL). After washes (3 x 5 minutes), sections were incubated in secondary antibodies Alexa Flour 488 (anti-mouse, 1:1000) for 6 hours and Alexa Flour 594 (anti-rabbit, 1:1000) for 3 hours at room temperature. Post incubation, sections underwent PB washes and then, sections were treated with Vector TrueVIEW Autofluorescence kit (SP-8400; Vector Labs, CA; 3 cases) for 2 minutes at room temperature to remove any background from aldehydes. After this step, tissue was washed 3 times with PB and subsequently, mounted in gelatin coated slides and coverslipped with VectaShield mounting medium with DAPI.

### Calretinin (CR^+^) and calbindin (CB^+^) DAB reactions

Brain sections that included midline thalamus were incubated for 48 hours at room temperature in rabbit calretinin (CR) primary antibody (1:2000, RPCA-Calret; Encore Bio, FL) OR mouse calbindin (CB) primary antibody (1:500, MCA-5A9; Encore Bio, FL) in 5mL of 0.1% BSA. After this period, tissue was washed 3 times for 5 minutes in 0.1 M PB and placed in goat anti-rabbit OR anti-mouse biotinylated secondary antibodies respectively (1:500, BP-9200/9100; Vector Labs, CA) for 6 hours. After PB washes, the tissue was reacted in a solution containing avidin–biotin complex (Vectastain Elite ABC Kit, PK-6100; Vector Laboratories, CA) for one hour at room temperature, followed by three 5-minute rinses in 0.1 M PB. The peroxidase reaction was produced by incubating the sections for 5 to 12 minutes in a DAB substrate solution (SK-4100; Vector Labs, CA). Reacted tissue was then mounted on gelatin coated slides, dehydrated in methanol and xylene before being cover-slipped with Permount or Vectashield antifade mounting medium with DAPI (H-1900, Vector Labs, CA).

### Imaging and data analysis

For each subject, sections were imaged at different rostro-caudal levels of midline thalamus (β range −1.08/−3.0). Cases with similar rostral, mid and caudal levels were grouped together for cell quantification across reactions, nuclei and levels. Schematic drawings (overlays) from Swanson Rat Brain Atlas (2018) were used to define the boundaries of the thalamic nuclei. The regions were quantified automatically (see below) to identify immunoreacted cell bodies at each of the selected brain levels. Atlas overlays were placed on top of original captures using Adobe Illustrator ^®^ (Adobe Systems Inc., San Jose, CA). Cell quantification was done on one hemisphere based on overall quality to avoid intrinsic confounds such as large blood vessels that might be present in any one section.

Immunofluorescence from PV^+^, CB^+^ and CR^+^ tissue, and retrogradely labeled RE neurons from mPFC and HC injections, was imaged using an Olympus FV1200 confocal microscope at 10X, 20X and 60X focusing on midline thalamic structures (PVT, PT, RH and RE) using standard filter cubes for red fluorescence (excitation 545nm, emission 605nm), green fluorescence (excitation 470nm, emission 525nm) and DAPI (excitation 350nm, emission 460nm). Captures at 60X magnification (oil immersed) were obtained to further explore and verify cell body staining. DAB peroxidase stained sections were captured using an Olympus BX51 brightfield microscope at 20X magnification.

### Automated cell counts

Quantification of neurons in all thalamic regions of interest (ROI) was performed using a customized automated pipeline built in CellProfiler Software version 3.1.9 (cellprofiler.org) for objective cell counts. FIJI ImageJ (Version 2.0.0; NIH; Schindelin et al., 2012) was used for preprocessing. The data was extracted from CellProfiler using customized Python code through Anaconda Software (Version 2-2.4.0). CellProfiler has been previously used and validated for both fluorescence and chromogenic localization by other research laboratories (McQuin et al., 2018; Tollemar et al., 2018). We further validated the accuracy of CellProfiler by comparing the counts from 26 ROIs between two experienced counters (manually) and CellProfiler. The counts from Experimenter 1 and Experimenter 2, and their combined average were significantly correlated with the results yielded by CellProfiler (CellProfiler v. Experimenter 1, *r* = .0.956; CellProfiler v. Experimenter 2, *r* = 0.941; CellProfiler v. Average, *r* = 0.958; all *p* < .001). A test of inter-rater reliability showed a very high degree of reliability (Cronbach’s α = 9.78) between the counters total average and CellProfiler’s results.

### Fluorescence-based PV^+^, CR^+^ and CB^+^ cell counts (single/dual)

A pipeline was created for the quantification of PV^+^, CB^+^, CR^+^ or DAPI fluorescence-based cell counts. *CorrectIlluminationCalculate* and *CorrectIlluminationApply* functions from CellProfiler were applied onto separated RGB images in order to correct any uneven lighting artifacts and further reduce noise. The three channels were aligned based on the signal intensity values using the CellProfiler *Align* function. A restricted range of diameters and a set of intensity values were determined for each channel allowing for proper cell identification. A mask of the identified cells was created for each RGB channel. Dual-labeled CR^+^/CB^+^ cells were quantified by relating the masks to each other Using CellProfiler’s *Relate Objects* function.

### DAB CR^+^ and CB^+^ cell counts

A separate CellProfiler pipeline was created to quantify the number of DAB stained CR^+^ and CB^+^ cells. ROIs were isolated from overlayed brightfield images using Adobe Photoshop^®^. Brightfield images were converted to a greyscale and inverted using the *ImageMath* function in CellProfiler. This process helps reduce noise, enhance cell features, and makes it easier to identify non-cells bodies artifacts, thus avoiding over-quantification. Quantification of cell bodies was determined with a set range of intensity and diameter values.

### Counting RE neurons with collaterals to mPFC and HC

We identified RE cell clusters that projected to both mPFC and HC with the FV1200 confocal microscope with using a z-stack (0.5 microns optical sections) to confirm dual-labeling. The same tissue was imaged with the Olympus BX41 brightfield-epifluorescence microscope for dual-fluorescence and DAB imaging similar to previous reports (Al-Mashhadi et al., 2015; Majercikova et al., 2012; Young et al., 2005). Individual captures were made at 20X magnification for the following channels: red (RE→PFC), green (RE→HC), blue (DAPI) and brightfield (DAB CR^+^ or DAB CB^+^ cells). Captures were made in two prominent dorsal and ventral regions within a 545μm x 390μm area that contained the dual-projecting clusters in RE. The corresponding brightfield image of CR or CB was transposed onto the merged fluorescent image using Adobe Photoshop^®^. A separate layer was created to mark the location of CR^+^ cells with white ‘+’ signs and CB^+^ cells with cyan ‘+’ signs. The flattened image was used to verify whether there was an overlap between DAB CR^+^ or DAB CB^+^ cells with immunofluorescence reacted neurons. Here, counts were performed manually by two experienced experimenters and averaged. DAPI cell counts were done using a custom CellProfiler pipeline.

### Cell soma (body) size

Using CR and CB dual reacted immunofluorescence sections, we measured soma size (μm^2^) in PVT and RE using FIJI Image J. We sampled cells in a 200μm x 200μm region of PVT and RE. Counts were made in medial, dorsolateral, and ventrolateral subregions of each nuclei. Cell bodies were outlined with the freehand tool in FIJI and measured using a set calibration scale from the microscope.

### Cell radius distance

In merged fluorescent and brightfield images, the radial distance of CR^+^ or CB^+^ cells from a small sample of dual mPFC-HC cells in RE was measured by imposing a 100μm radius circle centered on each individual dual labeled cell. Then, the distance from the center of dual labeled cells to all CR^+^ or CB^+^ cells within this area was measured using Image J.

### Statistical analysis

For dual reacted CB and CR immunofluorescence data, two-way repeated measures analysis of variance (ANOVAs) were performed to compare interaction or main effects of region (nuclei) and calcium binding protein expression type on cell area density across the rostro, medial and caudal levels, followed by one way-ANOVAs and multivariate/ pairwise comparisons (with Bonferroni correction) when the differences were statistically significant. For DAB reacted tissue, two-way ANOVAs were used to compare region (nuclei) and calcium binding protein expression type differences on cell area density. When significant, Bonferroni post hoc tests and pairwise comparisons followed. Effect size was performed using Hedges’d given the unequal sample sizes. Finally, a linear regression analysis was performed to assess the relationship between DAB cell counts and distance from dual projecting cells. All collected data was tallied and saved in Microsoft Excel and subsequently analyzed in SPSS (version 26). An alpha of 0.05 was considered statistically significant for all analysis.

## RESULTS

### Absence of PV^+^ cells in midline thalamus

We first looked for evidence of PV^+^ cell body labeling in the midline thalamus (PVT, PT, RE, and RH). We did not find PV^+^ cell body labeling in any of the three rostral-caudal sections selected to span the length of the midline thalamus (n = 4 rats, 12 coronal sections)(Fig. 1A-C). The lack of PV expression in midline thalamic cells stood in stark contrast to prominent PV^+^ cell body labeling in other brain regions with well-established PV^+^ cell populations including the thalamic reticular nucleus of the thalamus (TRN), the hippocampus, lateral and basolateral amygdala, striatum, and cortex (Fig. 1A, D-E). We also observed dense PV^+^ fibers and puncta throughout much of the nearby lateral thalamus and striatum. In TRN, we saw uniformly dense populations of PV^+^ cells with large immunonegative nuclei, which is characteristic of PV^+^ labeling in TRN neurons (Fig 1D;(Arai et al., 1994; Celio, 1990; Csillik et al., 2005; Kirichenko et al., 2017). Likewise, the distribution of PV^+^ cells in cortex was organized by layers and was comparatively more sparse (Fig 1E; also see Van Brederode et al., 1991; Ahn et al., 2017). Despite the absence of PV^+^ cell bodies in the midline thalamus, parvalbumin labeling was still abundant in the form of PV^+^ puncta that were most often clustered near or between cell bodies (Fig. 1B-C, insets). In some cases, PV^+^ puncta were observed enveloping entire cell bodies. More commonly, PV^+^ puncta formed asymmetrical cluster densities that were biased toward one pole and formed a rough spherical cap. Generally, the observations are in line with previous descriptions of PV^+^ labeling in midline thalamus (Celio, 1990; Arai et al., 1994).

**Figure 1.**
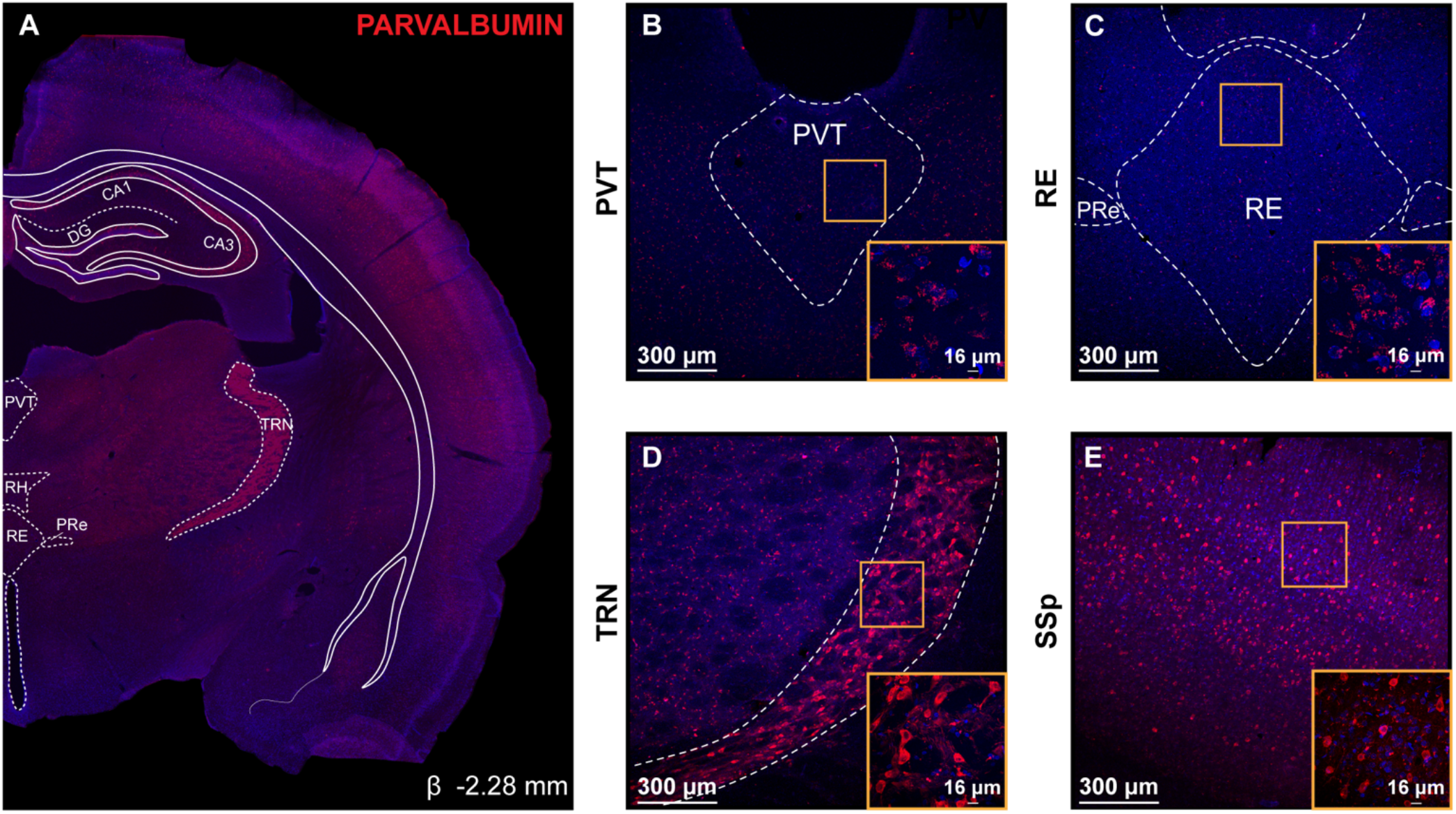
Absence of parvalbumin (PV^+^) cell bodies in midline thalamus. **A:** Representative coronal section (β −2.28mm) showing PV^+^ cell body expression throughout several regions of the brain. PV^+^ expression shown in red and DAPI in blue. Overlay shown adapted from Swanson (2018) to highlight thalamic structures. **B-E:** Confocal images showing PV immunoreactivity in PVT (**B**), RE (**C**), TRN (**D**) and SSp (**E**). Neither PVT nor RE contain PV^+^ cell bodies, however PV immunoreacted puncta was abundant near or between their cell bodies (**B-C** insets). Scale bar = 300μm. Inset scale bar= 16μm. **D:** PV^+^ cell bodies with a characteristic large immunonegative nuclei were seen in TRN (**D** inset). **E**: PV^+^ cell bodies were also observed in SSp cortex showing their distinct sparse but organized layer distribution (**E** inset). Gold squares represent regions of 60X magnification shown in inset. Abbreviations: β, bregma; CA1, CA1 subfield of the hippocampus; CA3, CA3 subfield of the hippocampus; DAPI, 4’,6-Diamidino-2-phenylindole dihydrochloride; DG, dentate gyrus; PV, parvalbumin; PVT, paraventricular; PRe, perireuniens; RE, nucleus reuniens; RH, rhomboid; SSp, primary somatosensory cortex; TRN, thalamic reticular nucleus.

### CR and CB expression in midline thalamus

Next we examined CR^+^ and CB^+^ labeling in three rostral-caudal sections because of known differences along this axis of the midline thalamus (Arai et al., 1994; Celio, 1990; Rogers & Résibois, 1992; Winsky et al., 1992). Each coronal section was immunofluorescence reacted for both CR and CB. A general finding was that CR^+^ and CB^+^ cell distributions show different clustering zones within and between PVT, PT, RE and RH. A clear overall pattern emerged in the CR^+^ and CB^+^ labeling that similarly organized the cytoarchitecture of the dorsal midline thalamus (PVT and PT) and ventral midline thalamus (RH and RE). That is, CR^+^ zones were dense medial-laterally against the dorsal and ventral 3V and ran dorso-ventrally along the midline away from their respective 3V forming the shape of a “T” or “Y” in dorsal midline thalamus, or a similar inverted pattern in ventral midline thalamus. In relation to CR^+^ cells, CB^+^ cells were clustered more ventro- or dorso-laterally (for dorsal and ventral midline thalamus, respectively). CR^+^ and CB^+^ zones partially overlapped and contained dual-labeled CR^+^/CB^+^ cells. Although these CR^+^ and CB^+^ patterns were similar in dorsal and ventral midline thalamus, there was generally more complexity to this organization in ventral midline thalamus.

### CR^+^ and CB^+^ labeling distributions in rostral midline thalamus

In rostral sections (β −1.44; n = 4 rats), CR^+^ cell and fibers densities were found in several regions including the hypothalamus, striatum, central and medial amygdala, medial divisions of thalamus, and cortex (Fig. 2A, magenta) notably including a very prominent CR^+^ fiber band in the superficial layers of entorhinal cortex (Wouterlood et al., 2001). In the midline thalamus, CR^+^ cells were predominantly located in the dorsal, medial and ventral portions (Fig. 2B & C, magenta). Whereas, CB^+^ cells and fibers often expressed in the same regions as CR^+^ cells, labeling was more prominent in the lateral portions of the thalamus, internal capsule, basolateral amygdala, and globus pallidus (Fig. 2A, green). In the midline thalamus, CB^+^ cells chiefly labeled the lateral portions (Fig. 2B & C). Generally, we observed that CR^+^ and CB^+^ cell distributions organized into well-defined zones within the midline thalamus giving the impression that there are important calcium binding protein specific subregions within PVT, PT and RE.

**Figure 2.**
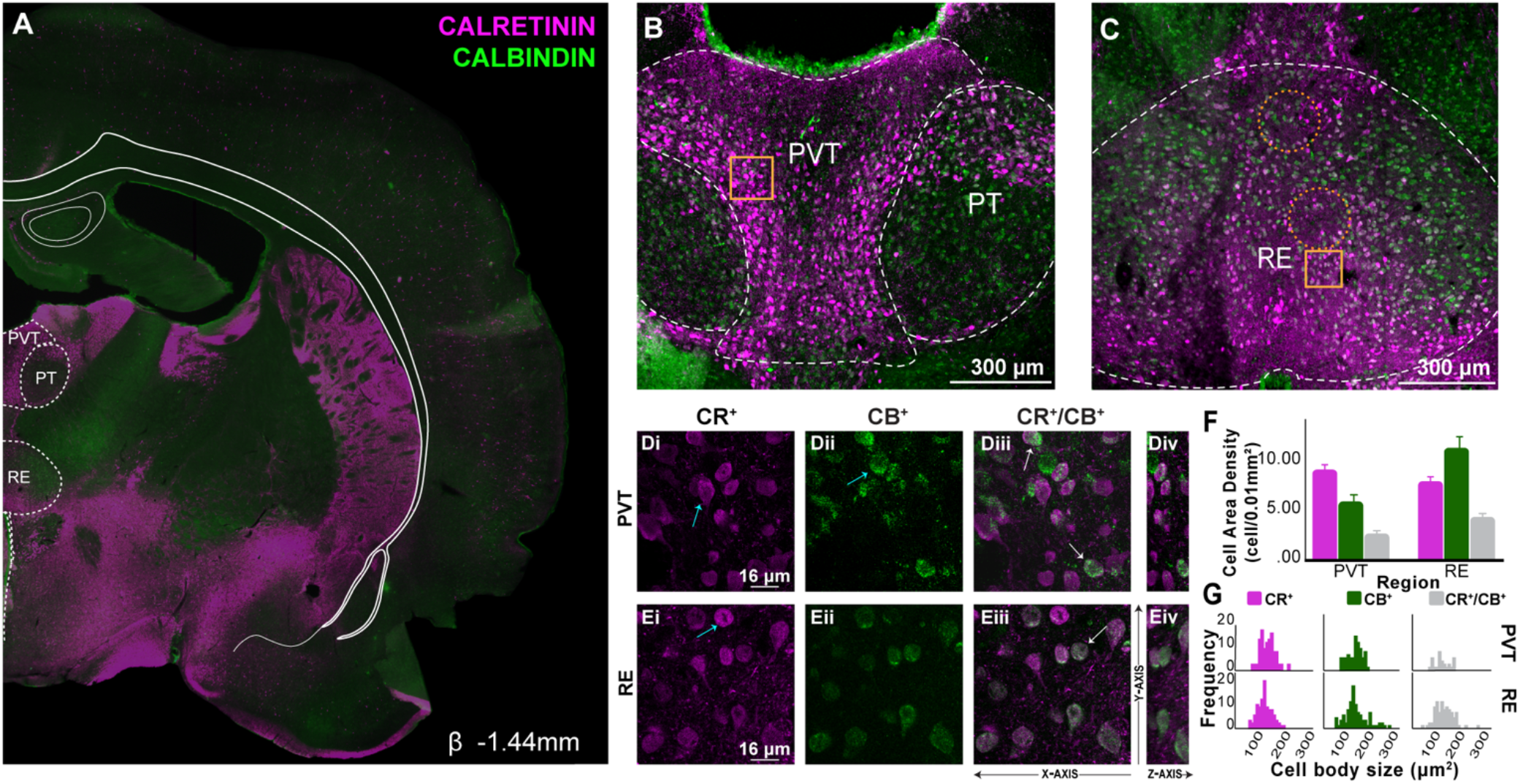
CR and CB labeling in rostral midline thalamus. **A:** Representative coronal section (β −1.44mm) showing immunofluorescent localization of CR^+^ and CB^+^ cell and fiber densities. Overlay shown adapted from Swanson (2018) to highlight midline thalamic structures. CR shown in magenta, CB in green. **B**: Confocal image demonstrating distribution of CR^+^ and CB^+^ cells in PVT and PT. In PVT, CR was prominent in dorsolateral and ventromedial regions, and CB ventrally in PT. **C**: A similar but inversed distribution was observed in RE where CR^+^ cells were prominent in ventrolateral and dorsomedial regions, while CB^+^ cells were prominent laterally. Gold squares represent region of 60X magnification shown in inset. Orange dotted circles indicate regions where calcium binding protein cell expression is sparse or absent. Scale bar = 300μm. **D:** Confocal images illustrating CR^+^ **(D_i_)**, CB^+^ **(D_ii_)** and dual labeled CR^+^/CB^+^ **(D_iii_)** immunoreacted cell bodies in PVT. Three distinct cell populations were identified: CR^+^ only cells (**D_i_**, blue arrow), CB^+^ only cells (**D_ii_**, blue arrow) and dual CR^+^/CB^+^ cell bodies (**D_iii_**, white arrows). The Z-axis from these optical sections are shown to the right **(D_iv_)**. E: Confocal images in RE **(E_i_-E_iv_)**. Scale bar = 16μm. **F:** Comparison of CR^+^ and CB^+^ cell area density in PVT and RE (cells/0.01mm^2^) in rostral levels. Error bars represent SEM. **G:** Frequency distribution of CR^+^, CB^+^ and CR^+^/CB^+^ immunoreacted cell body size (μm^2^) in rostral PVT and RE. Abbreviations: β, bregma; CB, calbindin; CR, calretinin; PVT, paraventricular; PT, paratenial; RE, nucleus reuniens, SEM, standard error of the mean.

### Distinctive CR^+^ and CB^+^ labeling in rostral PVT and PT

Next, we focused on CR^+^ and CB^+^ labeling in the dorsal midline thalamic nuclei (PVT and PT; Fig. 2B). While PVT and PT are often considered together, their CR^+^ and CB^+^ cell distributions indicated that they contain different functional zones. In PVT, CR^+^ cells were bright, large, and clustered together in a chain formation that ran dorsoventrally and hugged the lateral borders (*M* = 9.168, *SD* = 1.146; density in cells per 0.01mm^2^; Fig. 2B & D_i_). By contrast, few CR^+^ cells were visible in dorsal and medial PVT, although these areas were heavily populated with CR^+^ puncta. CB^+^ cells stained much lighter, but exhibited similar topography within PVT (*M* = 5.928, *SD* = 1.490; density in cells per 0.01mm^2^; Fig. 2B & D_ii_). Dual-labeled CR^+^ and CB^+^ cells were most prominent in the lateral portions of PVT (Fig. 2C & D_iii_). These dual-labeled CR^+^/CB^+^ cells accounted for 29.22% of CR^+^ cells and 45.19% of CB^+^ cells in PVT (Fig. 2F).

Different from PVT, PT had a clear segregation between CB^+^ and CR^+^ cell topographies (Fig. 2B). In PT, CR^+^ cells (*M* = 3.600, *SD* = 1.978; density in cells per 0.01mm^2^) were practically absent from ventral PT but were abundant in dorsal PT. This CR^+^ and CB^+^ labeling delineates rostral PT into dorsal and ventral subregions (Fig 2B). CB^+^ cells (*M* = 8.845, *SD* = 4.847; density in cells per 0.01mm^2^) were observed throughout PT, usually intermingled or overlapping with CR^+^ cells in dorsal PT, or as an independent population in ventral PT. Dual-labeled CR^+^/CB^+^ cells (*M* = 2.398, *SD* = 2.086; density in cells per 0.01mm^2^) were located in dorsal PT. CR^+^/CB^+^ cells accounted for 66.67% of CR^+^ cells and 27.12% of CB^+^ cells in PT.

### Rostral RE exhibits CR^+^ and CB^+^ defined zones

In rostral sections, the ventral midline thalamus is composed entirely of RE, which showed distinctive patterns of CR^+^ and CB^+^ labeling (Fig. 2C). Rostral RE CR^+^ cells were bright, large, and their intensity increased from dorsal to ventral borders (*M* = 7.990, *SD* = 1.042; density in cells per 0.01mm^2^). Generally, CR^+^ cells in rostral RE were concentrated in dorsal, middle, and ventral regions and practically avoided lateral areas, with the exception of a few cells that dual-labeled with CB^+^ cells (Fig. 2C & E). RE CB^+^ cells (*M* = 11.383, *SD* = 2.404; density in cells per 0.01mm^2^) were present throughout the whole body of RE but were more densely packed and separated from CR^+^ cells in dorsolateral regions. In dorsal and medial RE, CR^+^ and CB^+^ cells were loosely distributed throughout and cells were visibly smaller in size compared to cells in the dorsolateral regions (Fig. 2C, E_i_-E_ii_; Table S1). Patches of dual-labeled CR^+^/CB^+^ cells (*M* = 4.365, *SD* = 0.823; density in cells per 0.01mm^2^) were common in dorsal, medial and ventral subdivisions of RE (Fig. 2C & E_iii_). These dual-labeled CR^+^/CB^+^ cells accounted for 54.63% of CR^+^ cells, and 38.35% of CB^+^ cells in RE. A prominent region of CR^+^/CB^+^ cells was located in the boundary between RE mid-ventral regions and RE lateral areas (Fig. 2C). We also noted there were visible circular bands of CR^+^ and CB^+^ cells that contained a sparsely labeled center (Fig. 2C, dotted circles), although DAPI indicated the presence of cells within these circles. These results show CR^+^, CB^+^, and CR^+^/CB^+^ zones that easily delineate rostral RE.

### PVT and RE show opposing CR^+^, CB^+^ and CR^+^/CB^+^ cell densities and cell size patterns in rostral RE

Next we focused on comparing PVT and RE, the largest and most studied regions of midline thalamus, which are well known for their different functional roles (Cassel et al., 2013; Hsu et al., 2014; Kawano, 2001; Matzeu et al., 2014; Vertes et al., 2015). The overall CR^+^ and CB^+^ cell labeling patterns in rostral midline thalamus suggested clear differences between PVT and RE with more CR^+^ cells in PVT and more CB^+^ cells and dual-labeled CR^+^/CB^+^ cells in RE. First, we compared CR^+^ and CB^+^ cell densities in PVT and RE. There was a significant effect of region (*F*_(1,3)_ = 32.773, *p* = 0.011) such that CB^+^ cell density in RE was slightly more than PVT, but no main effect of cell densities by calcium binding protein expressed (*F*_(1,3)_ = 0.004, *p* = 0.955), and a significant interaction effect (*F*_(1,3)_ = 4.907, *p* = 0.006), indicating opposite calcium binding expression proportions between PVT and RE (Fig. 2F). There were also significantly higher densities of dual-labeled CR^+^/CB^+^ cells in RE than PVT (1.63:1, RE:PVT; paired-samples *t*_(3)_ = 9.268, p = 0.003). A comparison of CR^+^ and CB^+^ cell sizes in PVT and RE (Fig. 2G) showed no significant main effect of cell size by calcium binding protein expressed (*F*_(1,324)_ = 0.467, *p* = 0.495) or region (*F*_(1,324)_ = 0.633, *p* = 0.427), but there was a significant interaction effect (*F*_(1,324)_ = 5.404, *p* = 0.021). In PVT, CR^+^ cells tended to be larger compared to RE, and CB^+^ cells were larger in RE compared to PVT (Fig. 2G). Further in PVT, cells were largest laterally and smallest medially. In RE, CB^+^ cells were largest in dorsolateral subdivisions, but smaller in medial and ventrolateral divisions. Dual-labeled CR^+^/CB^+^ cells in PVT and RE were not significantly different (paired-samples *t*_(105)_ = 0.411, *p* = 0.523). Detailed cell size measurements are provided in supplemental table S1.

### CR^+^ and CB^+^ labeling in mid-levels of the midline thalamus

In mid-level sections (β −1.78; n = 5 rats), the overall CR^+^ (Fig. 3A, magenta) and CB^+^ (Fig. 3A, green) cell and fiber densities were similarly distributed to the rostral levels. A notable change was more abundant CR^+^ labeling in habenula, mediodorsal, anterodorsal and centromedial nuclei of the thalamus, and CB^+^ cell densities in interanteromedial and anteriomedial nuclei of the thalamus, and the hypothalamus. CR^+^ cells were visually more prominent in dorsal midline thalamus and CB^+^ cells were more prominent in ventral midline thalamus, but CR^+^ and CB^+^ cells were found in all regions including PVT, PT, RE and RH to differing degrees (Fig. 3B & C).

**Figure 3.**
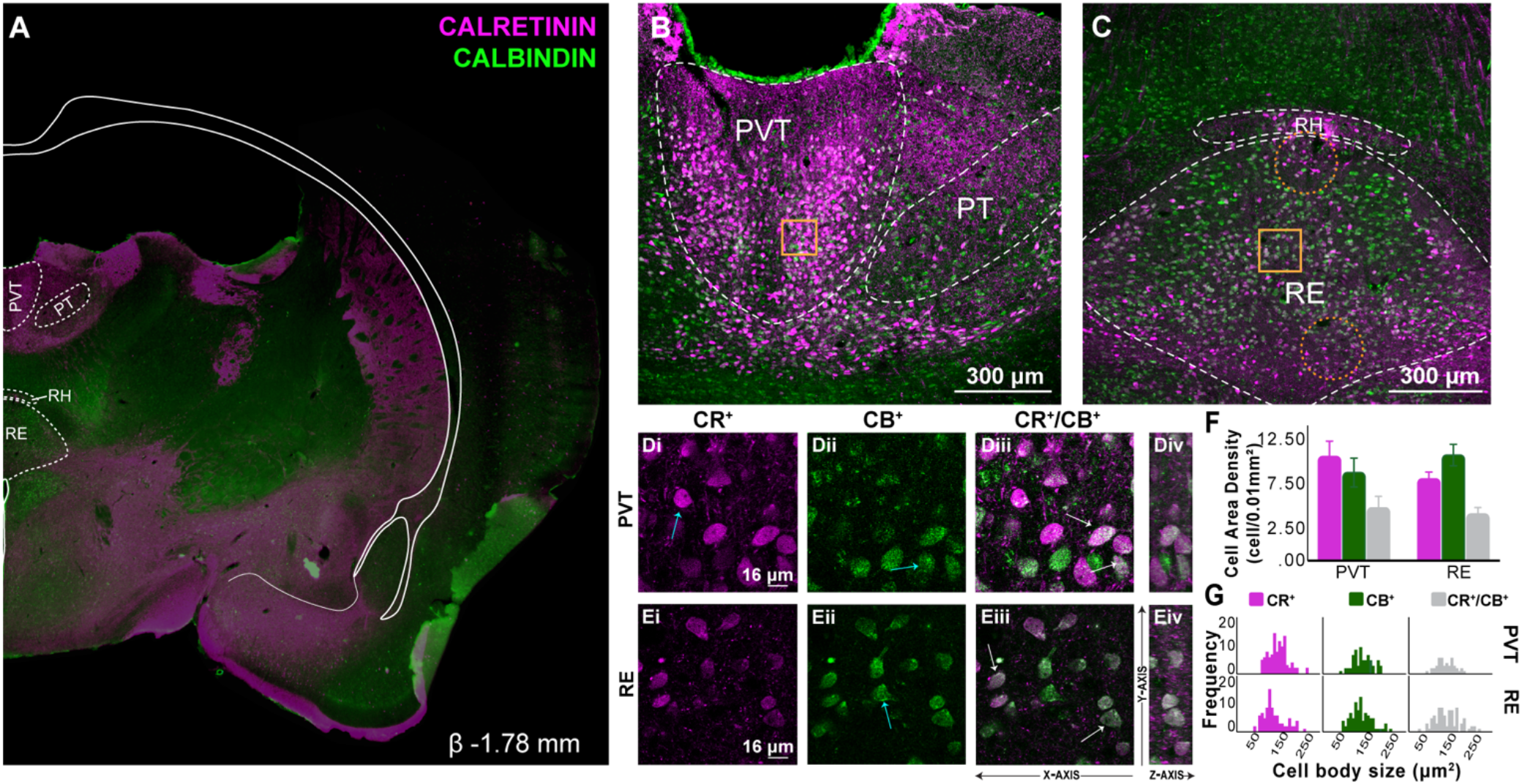
CR and CB labeling in mid levels of midline thalamus. **A:** Representative coronal section (β −1.78 mm) showing immunofluorescent localization of CR^+^ and CB^+^ cell and fiber densities in mid-levels of the rostro-caudal axis of the thalamus. Overlay shown adapted from Swanson (2018) to highlight midline thalamic structures. CR shown in magenta, CB in green. Fi, RSD and motor-sensory cortex layers missing from tissue section. **B**: Confocal image demonstrating distribution of CR^+^ and CB^+^ in PVT and PT. Compared to rostral levels, PVT CR^+^ and CB^+^ cells re-distributed more ventral and laterally. CR^+^ fibers were abundant in dorsal PVT and PT (below 3V). **C**: In RE, a similar but inverse re-distribution was also observed, with CR^+^ cells prominent ventral and laterally and CB^+^ cells in dorsolateral regions. A zone of CR^+^ fibers emerged ventromedially just above 3V. Gold squares represent regions of 60X magnification shown in inset. Orange dotted circles indicate regions in which there is sparse or no expression of calcium binding cells. Scale bar = 300μm. **D:** Confocal images illustrating CR^+^ **(D_i_)**, CB^+^ **(D_ii_)** and dual labeled CR^+^/CB^+^ **(D_iii_)** immunoreacted cell bodies in PVT. At this level, the same three calcium binding cell types were visualized: CR^+^ only cells (**D_i_**, blue arrow), CB^+^ only cells (**D_ii_** blue arrow) and dual CR^+^/CB^+^ cell bodies (**D_iii_**, white arrows). The Z-axis from these optical sections are shown to the right **(D_iv_)**. **E:** Confocal images in RE **(E_i_-E_iv_)**. Scale bar = 16μm. **F:** Comparison of CR^+^ and CB^+^ cell area density in PVT and RE in mid-levels of midline thalamus (cells/0.01mm^2^). **G:** Frequency distribution of CR^+^, CB^+^ and CR^+^/CB^+^ immunoreacted cell body size (μm^2^) in PVT and RE at mid-levels of the thalamus. Error bars represent SEM. Abbreviations: β, bregma; CB, calbindin; CR, calretinin; PVT, paraventricular; PT, paratenial; RE, nucleus reuniens; RH, rhomboid; Fi, fimbria of the hippocampus, RSD, retrosplenial dysgranular cortex, SEM, standard error of the mean; 3V, third ventricle.

### Distinctive CR^+^ and CB^+^ cell labeling in PVT and PT at mid-levels of midline thalamus

In mid-level coronal sections, PVT CR^+^ cells were dense (*M* = 11.346, *SD* = 3.771, density in cells per 0.01mm^2^), bright, and packed together in large circular clusters located laterally (Fig. 3B). There was notably sparse cell labeling in dorsal PVT with extensive CR^+^ puncta organized in dense fiber fields that often ran along the dorsal-ventral axis and thickened at the 3V border. CB^+^ cells were less dense (*M* = 9.604, *SD* = 3.516, density in cells per 0.01mm^2^), appeared lighter than CR^+^ cells, and largely overlapped topographically with CR^+^ cells (Fig. 3B, D_i_-D_iv_). Unlike CR^+^ cells, CB^+^ cells showed no clear tendency to cluster. Dual-labeled CR^+^/ CB^+^ cells were found interspersed equally in CR or CB cell rich areas and accounted for 50.56% of CR^+^ cells and 62.84% of CB^+^ cells.

In contrast to rostral levels, mid-level PT CR^+^ cells were mostly confined to ventral areas and comingled with CB^+^ cells (Fig. 3B). However, dorsal PT had profuse CR^+^ puncta visibly organized into fibers traversing around cells in meshed-wire pattern (e.g., Moyer et al., 2011). Generally, CR^+^ cells were very sparse (*M* = 1.578, *SD* = 0.826, density in cells per 0.01mm^2^) and CB^+^ cells were abundant (*M* = 9.805, *SD* = 3.935, density in cells per 0.01mm^2^). Dual-labeled CR^+^/CB^+^ cells were also sparse accounting for 48.65% of CR^+^ cells and 7.83% of CB^+^ cells.

### Distinctive CR^+^ and CB^+^ labeling in RE and RH at mid-levels of midline thalamus

In mid-level sections, RE and RH showed distinctive patterns of CR^+^ and CB^+^ labeling (Fig. 3C). Compared to rostral levels, mid-level RE CR^+^ cells were bright and abundant (*M* = 8.910, *SD* = 1.683, cells per 0.01mm^2^) with a slight shift in location ventrally and laterally towards the early formation of RE wings, and defined the lower border. CR^+^ cells were also seen along the lateral borders of RE, and dorsomedially (Fig. 3C). By comparison, CB^+^ cells (*M* = 11.520 *SD* = 2.622, cells per 0.01mm^2^), were localized throughout RE with independent (non-overlapping) populations in dorsal and dorsolateral regions. Notably at this level, RE CB^+^ cells in dorsolateral portions showed a very large cell body size that contrasted to that of smaller RE CB^+^ cells in its ventrolateral division (Fig. 3E_i_-E_ii_ & Table S1). Dual-labeled CR^+^/CB^+^ were mostly located along the lateral and ventral borders of RE, and as patchy clusters in centromedial and ventrolateral divisions of RE (Fig. 3C). CR^+^/CB^+^ cells accounted for 55.05% of CR^+^ cells and 41.73% of CB^+^ cells (Fig. 3E_iii_-E_iv_).

In RH, CR^+^ cells were scarce (*M* = 1.760, *SD* = 1.017, cells per 0.01mm^2^), and presented similar brightness and circular organization of that of PVT cells. CR^+^ cells were primarily confined to medial RH (Fig. 3C), with CR^+^ fibers present laterally. By contrast, CB^+^ cells were not bright but were plentiful throughout RH (*M* = 10.373, *SD* = 2.703, cells per 0.01mm^2^) with a tendency to cluster in lateral ends. Dual-labeled CR^+^/CB^+^ cells accounted for 50.71% of CR^+^ cells and 8.60% of CB^+^ cells.

### Mid-level PVT and RE show opposing CR^+^, CB^+^, and CR^+^/CB^+^ cell density and cell size patterns

In mid-levels, PVT and RE differences remained clear with PVT showing dominant CR^+^ labeling with CB^+^ cells interspersed, and RE showing more CB^+^ labeling dorsally and medially with CR^+^ cell zones seen ventrally and laterally. We compared overall CR^+^ and CB^+^ cell densities in PVT and RE. There was no main effect of cell density by calcium binding protein expressed (*F*_(1,4)_ = 0.317, *p* = 0.604) or region (*F*_(1,4)_ = 0.012, *p* = 0.918), but there was a significant interaction effect (*F*_(1,4)_ = 8.702, *p* = 0.042). As in rostral sections, mid-level PVT had relatively more CR^+^ cells, and RE had more CB^+^ cells (Fig. 3F). Unlike in rostral sections, there were no significant differences in the densities of dual-labeled CR^+^/CB^+^ cells (paired-samples *t*_(4)_ = 0.366, *p* = 0.733). A comparison of CR^+^ and CB^+^ cell sizes in PVT and RE (Fig. 3G) showed no differences by calcium binding protein expression (*F*_(1,335)_ = 0.040, *p* = 0.843), a significant effect of region (PVT; *F*_(1, 335)_ = 6.406, *p* = 0.012), and no significant interaction effect (*F*_(1,335)_ = 0.994, *p* = 0.320). That is, PVT cells tended to be slightly larger (8.33%) than RE (Fig. 3G & Table S1).

### CR^+^ and CB^+^ labeling distributions in caudal midline thalamus

In caudal sections (β-2.76; n = 4 rats), there were some notable variations in the overall distribution of CR^+^ and CB^+^ labeling (Fig. 4A) including that CR^+^ cell and fiber densities were now seen in basolateral amygdala, and CB^+^ staining increased in prominence in striatum. In midline thalamus, CR^+^ staining dorsally remained high (Fig. 4B). The balance of CR^+^ and CB^+^ labeling in ventral midline thalamus shifted whereby RE CR^+^ labeling intensity increased, and RH CB^+^ staining intensity increased, as compared to more rostral sections (Fig. 4C). Nuclei located centromedially in thalamus had numerous CB^+^ cells, while CR^+^ cells were less abundant (Fig. 4A). As before, cells expressing both types of calcium binding proteins were found in all midline thalamic structures examined in detail including PVT, RE, and RH.

**Figure 4.**
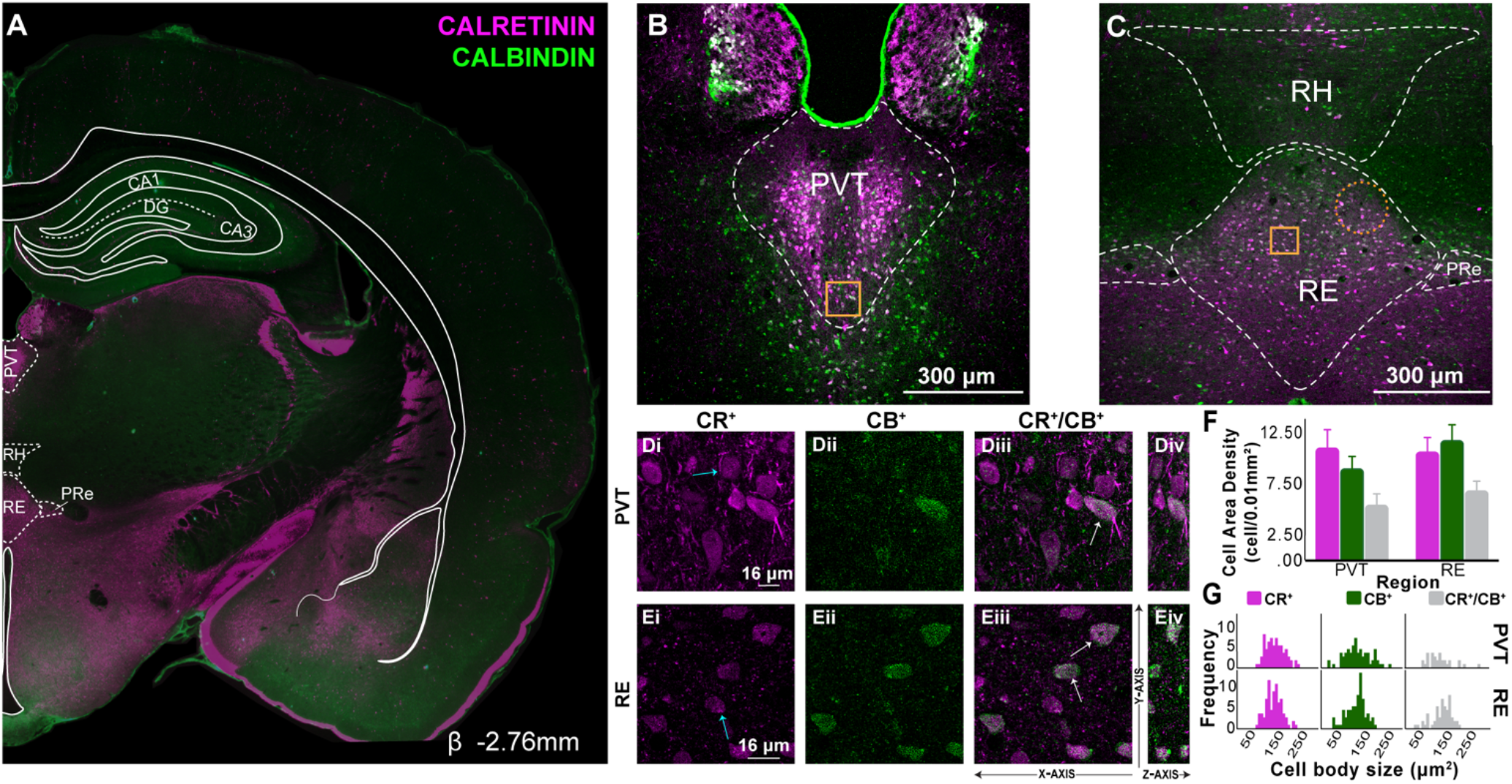
CR and CB labeling in caudal midline thalamus. **A:** Representative coronal section (β −2.76 mm) showing immunofluorescent localization of CR^+^ and CB^+^ cell and fiber densities in caudal midline thalamus. Overlay shown adapted from Swanson (2018) to highlight midline thalamic structures. CR shown in magenta, CB in green. **B**: Confocal image demonstrating distribution of CR^+^ and CB^+^ cells in PVT and PT caudally. In PVT, CR^+^ cells clustered mediolaterally and CB^+^ cells were predominantly seen in PVT ventral borders. **C**: In RE, CB^+^ cells were prominent in dorsal and lateral borders, while CR^+^ cells were observed more mediolaterally, often overlapping with CB^+^ cells. CR^+^ fibers continued to be abundant ventrally (above 3V). Gold squares represent region of 60X magnification shown in inset. Orange dotted circles indicate regions in which there is sparse or no expression of calcium binding cells. Scale bar = 300μm. **D:** Confocal images illustrating CR^+^ **(D_i_)**, CB^+^ **(D_ii_)** and dual labeled CR^+^/CB^+^ **(D_iii_)** immunoreacted cell bodies in PVT. As in previous levels, three cell types are visualized: CR^+^ only cells (**D_i_**, blue arrow), CB^+^ only cells (not shown) and dual CR^+^/CB^+^ cell bodies (**D_iii_**, white arrows). The Z-axis from these optical sections are shown to the right **(D_iv_)**. **E:** Confocal images in RE **(E_i_-E_iv_)**. Scale bar = 16μm. **F:** Comparison of CR^+^ and CB^+^ cell area density in PVT and RE in caudal levels of the midline thalamus (cells/0.01mm^2^). Error bars represent SEM. **G:** Frequency distribution of CR^+^, CB^+^ and CR^+^/CB^+^ immunoreacted cell body size (μm^2^) in caudal PVT and RE. Abbreviations: β, bregma; CA1, CA1 subfield of the hippocampus; CA3, CA3 subfield of the hippocampus; CB, calbindin; CR, calretinin; DG, dentate gyrus; PVT, paraventricular; PRe, perireuniens; RE, nucleus reuniens, RH, rhomboid, SEM, standard error of the mean.

### Caudal PVT CR^+^ and CB^+^ labeling

In caudal PVT, expression of CR^+^ cells (*M* = 10.950, *SD* = 3.588, cells per 0.01mm^2^) were bright and clustered tightly just off the midline running along the dorsal/ventral axis, similar to more rostral sections (Fig. 4B). Dorsal PVT was absent of any cell labeling for CR or CB, but was dense with CR^+^ fibers. CB^+^ cells (*M* = 8.935, *SD* = 2.412, cells per 0.01mm^2^) in caudal PVT lightly stained and were seen throughout the structure. CB^+^ cells were especially noticeable in the lateral and ventral portions (Fig 4B). Dual-labeling CR^+^/CB^+^ cells were prominent in dorsolateral areas accounting for 52.64% of CR^+^ cells and 56.93% of CB^+^ cells (Fig. 4Di-Div).

### Distinctive CR^+^ and CB^+^ labeling in RE, PRe, and RH in caudal midline thalamus

In caudal sections, distinct topographical organizations of CR^+^ and CB^+^ cells emerged in RH, RE, and PRe. Caudal RE CR^+^ cells (*M* = 10.560, *SD* = 2.845, cells per 0.01mm^2^) retained a bright color and were mostly located dorsolaterally, unlike earlier rostro-caudal levels. The ventral portion of RE was largely void of cells that labeled for either calcium binding protein. A network of CR^+^ fibers located closed to 3V, occupied most of this region along with a few CR^+^ cells located laterally. (Fig. 4C). CR^+^ cell expression was low in PRe (Fig. 4C), a structure known to be rich in RE neurons projecting to mPFC (Cassel et al., 2013; Dolleman-van der Weel et al., 2019). RE CB^+^ cells (*M* = 11.673, *SD* = 3.106), cells per 0.01mm^2^) were seen prominently along the lateral borders, where they overlapped heavily with CR^+^ cells (Fig. 4E_i_-E_iv_). CB^+^ cells were abundant throughout PRe. In the ventralmedial portion of RE, CB^+^ cells were few, small, and scattered (Fig. 4E; see Table S1). Dual-labeling CR^+^/CB^+^ cells were seen throughout the dorsal and medial subdivisions of RE and in PRe. CR^+^/CB^+^ cells accounted for 64.30% of CR^+^ cells and 58.18% of CB^+^ cells in RE (Fig. 4E).

In caudal RH, CR^+^ were largely absent (*M* = 2.163, *SD* = 1.368, cells per 0.01mm^2^), while CB^+^ cells (*M* = 7.630, *SD* = 5.344, cells per 0.01mm^2^) were abundant and present through the region (Fig. 4C). RH CR^+^ cells were mainly located in the mediodorsal and medioventral borders of RH, while CB^+^ cells were heavily distributed in the lateral wings with a relatively large size (Fig. 4C). Dual-labeled CR^+^/CB^+^ expression accounted for 31.02% of CR^+^ cells and 8.78% of CB^+^ cells.

### Caudal PVT and RE have different CR^+^, CB^+^, and CR^+^/CB^+^ cell density and cell size patterns

In caudal midline thalamus, the overall pattern of CR^+^ and CB^+^ cell densities in PVT and RE appeared to match well with more rostral sections. There were proportionally more CR^+^ than CB^+^ cells in PVT, and more CB^+^ than CR^+^ cells in RE, although these differences were moderate (Fig. 4F). We compared overall CR^+^ and CB^+^ cell densities in caudal PVT and RE. There was no main effect of calcium binding protein expression (*F*_(1,3)_ = 0.542, *p* = 0.515), or region (*F*_(1,3)_ = 0.519, *p* = 0.523), but there was a significant trend towards and interaction effect (*F*_(1,3)_ = 7.836, *p* = 0.68). Similar to mid-level sections, there was no significant difference in the proportion of dual-labeled CR^+^/CB^+^ cells in PVT and RE (paired-sample *t*_(3)_ = −1.258, *p* = 0.297). A comparison of CR^+^ and CB^+^ cell sizes in caudal PVT and caudal RE (Fig. 4G) showed no differences by calcium binding proteins (*F*_(1,310)_ = 1.996, *p* = 0.159), no region effect (*F*_(1,310)_ = 1.265, *p* = 0.262), and no interaction effect (*F*_(1,310)_ = 0.011, *p* = 0.917)(Fig. 4G).

### Distributions of CR^+^ and CB^+^ in RE subregions using 3,3’-Diaminobenzidine (DAB)

Notably, the results of the immunofluorescence experiments revealed topographically biased clusters of CR^+^ and CB^+^ cell populations in RE, the largest of the midline thalamic nuclei (Arai et al., 1994; Bokor et al., 2002; Winsky et al., 1992) that varied across the rostral-caudal axis. We further investigated the distribution of RE CR^+^ and CB^+^ cells with DAB in all the RE internal subdivisions from those established in Swanson brain atlas (2018) including five rostro-caudal levels of the rat’s midline thalamus (n = 13 rats, 7 CR and 6 CB) to confirm these patterns and densities. The results were entirely consistent with the immunofluorescence in that all subregions of RE tended to have higher CB^+^ cell area densities that CR^+^ cell densities, except RE medial (Fig. 5). The cell area densities found with the DAB staining of CR^+^ and CB^+^ was almost identical to that of our immunofluorescence reactions (see above and Table S2), and the topographies were the same. Figure 5C summarizes our findings by atlas subregion showing that CB^+^ cell densities were consistently higher than CR^+^ cell densities, with the exception of cells at RE medial (REm) division. The same was true of RE caudoposterial (REcp) division, but only at mediocaudal levels. Notably, in rostromedial (reuniens lateral-REl and reuniens ventral division-REv) and caudal levels (reuniens caudal dorsal division-REcd) CB^+^ cell densities were about twice the density of CR^+^ cells (with the exception of REm). We calculated the overall effect size across all RE subregions and levels and found a modest effect of CB^+^ density (Hedge’s d = .32).

**Figure 5.**
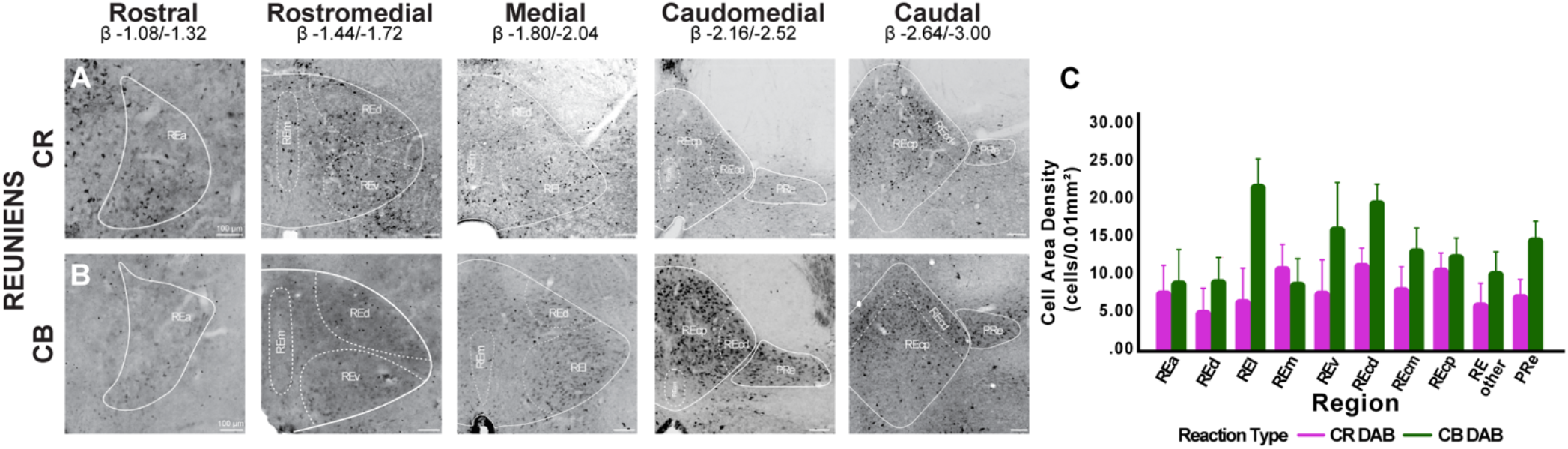
Distribution of DAB CR^+^ and DAB CB^+^ cells is not the same across all RE internal subdivisions. **A:** Brightfield images showing the distribution of DAB CR^+^ cells in RE across the rostro-caudal axis of the thalamus. CR^+^ cell area density varied depending on the subdivision of RE in which they were located. **B:** Distribution of DAB CB^+^ cells. When compared, CR^+^ and CB^+^ cell distribution in all RE’s subregions and across the rostral to caudal levels did not appear to be the same. Overlay shown adapted from Swanson (2018) to highlight all RE internal subdivisions. Scale bar = 100μm. **C:** Comparison of DAB CR^+^ and DAB CB^+^ cell area density (cells/0.01mm^2^) in all subdivisions of RE across the rostro-caudal axis. CB^+^ cell densities were higher than CR^+^ cell densities (except REm). Additionally, REl, REv and REcd subdivisions exhibited large CB^+^ cell densities compared to other RE subregions. A moderate size effect was found for CB^+^ cell area density across all levels and RE subdivisions (Hedges’ d=0.32). Abbreviations: β, bregma; CB, calbindin; CR, calretinin; DAB, 3,3’-Diaminobenzidine; PRe, perireuniens, RE, nucleus reuniens of the thalamus, REa, reuniens rostral division anterior part; REd, reuniens rostral division dorsal part; REl, reuniens rostral division lateral part; REm, reuniens rostral division median part; REv, reuniens rostral division ventral part; REcm, reuniens caudal division median part; REcd, reuniens caudal division dorsal part; REcp, reuniens caudal division posterior part.

### Dual-site mPFC-HC projecting RE neurons are not CB^+^ or CR^+^

A noteworthy feature of the midline thalamus is the presence of cells with monosynaptic projections to both mPFC and HC (Hoover & Vertes, 2012; Varela et al., 2014) which are thought to contribute to rhythmic synchrony and communication in the mPFC-HC memory system (M. J. Dolleman-van der Weel et al., 2019). Thus, we performed additional experiments to examine the calcium binding protein expression in dual mPFC-HC projecting cells in RE. To do this, we injected two AAV retrograde viral vectors in mPFC (prelimbic and infralimbic cortex; rAAV-CAG-tdTomato) and HC (ventral CA1; rAAV-CAG-GFP) and then imaged the resultant tdTomato (red) and GFP (green) expression in RE (Fig. 6A) (n = 4 rats). Coronal sections were counterstained with DAB to label CR (n = 2) and CB (n = 2) (see Figure S1). We found cells in RE that were dual-labeled (Fig. 6B, yellow) that clustered predominantly in consistent dorsal and ventral locations near the midline (Fig. 6B, cyan squares). Dual mPFC-HC projecting cells represented only a small proportion of RE cells in the area (Fig. 6C) consistent with previous reports (Hoover & Vertes, 2012; Varela et al., 2014). Next, we looked at whether CR^+^ or CB^+^ cells co-localized with dual mPFC-HC projecting cells in RE. Surprisingly, no CR^+^ and CB^+^ cells overlapped RE dual-projecting cells. We also noticed (as before in Figs. 2–4C) that there were CR^+^ and CB^+^ sparse areas, and interestingly these appeared to contain dual mPFC-HC projecting cells. To examine this visual impression quantitatively, we measured the distance from a sample of RE dual-projecting cells (n = 49) to every CR^+^ and CB^+^ cell within a 100-micron radius (D_ii_-E_ii_). There were very few CR^+^ and CB^+^ cells nearby RE dual-projecting cells, and this count progressively increased with distance (Fig. 6F_ii_-G_ii_). To confirm this relationship, we ran a linear regression between the cell counts by distance. We found that CR^+^ cell and CB^+^ cell counts both increased significantly as a function of distance from dual mPFC-HC projecting cells in RE (dorsal clusters: CR^+^ cells r = .57, *r^2^* = .328; CB^+^ r = .80 *r^2^* = .646; ventral clusters: CR^+^ cells r = .84, *r^2^* = .706; CB^+^ r = .84, *r^2^* = .888; all p’s < 0.01). Overall, these findings show that dual mPFC-HC projecting cells in RE are a neurobiologically unique cell type in that they lack CR and CB (and PV) expression, and that their local cytoarchitecture clusters them within CB^+^ and CR^+^ cell nests that occurred in multiple topographic locations within RE.

**Figure 6.**
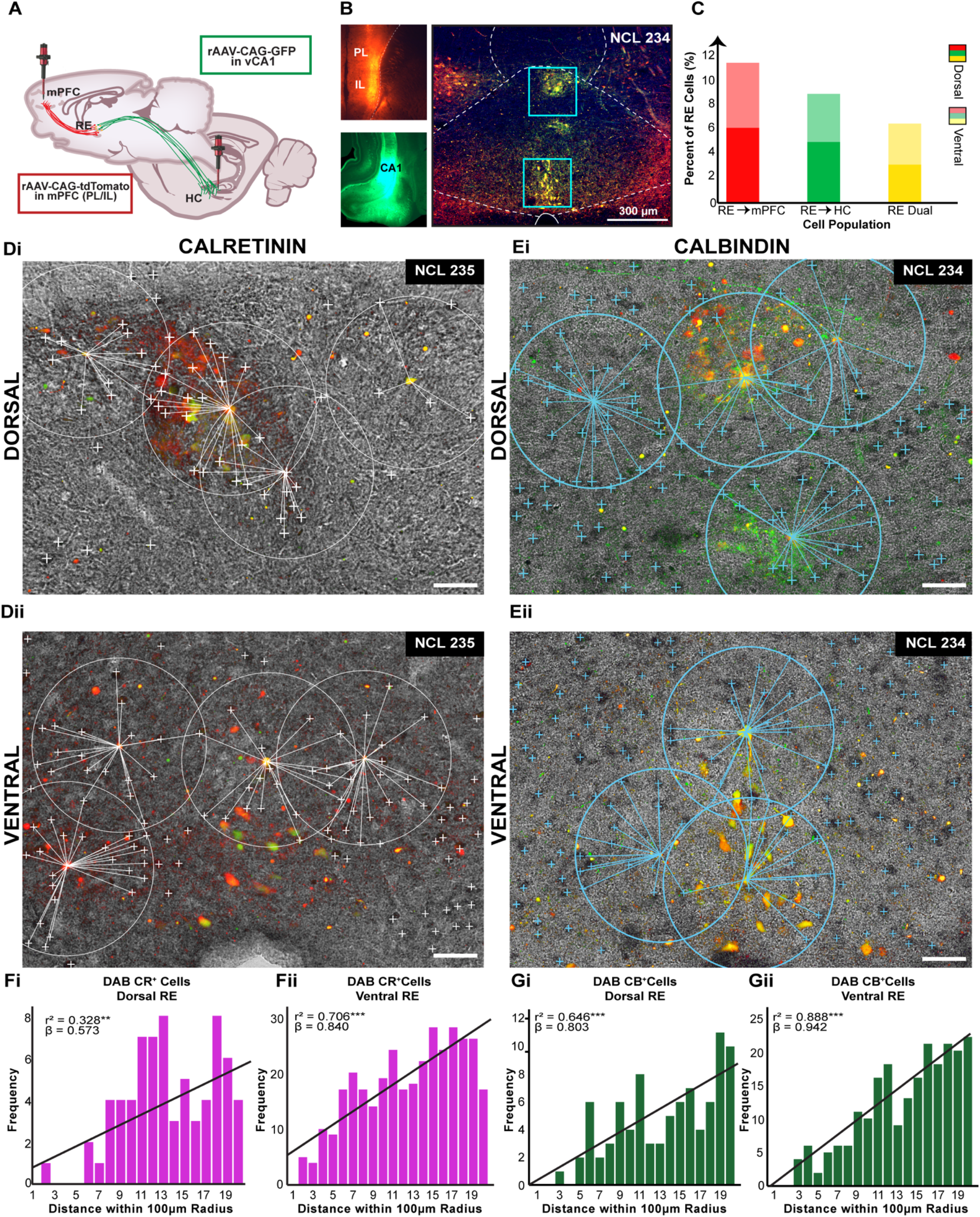
Dual-site mPFC-HC RE projecting cells are not CB^+^ or CR^+^ cells. **A**: Experimental design. Paired bilateral injections of rAAV-CAG-tdTomato and rAAV-CAG-GFP delivered in mPFC and HC. **B**: Injections’ spread was confined to PL/IL layer V/VII (top left) and vCA1 (bottom left). Retrogradely dual mPFC-HC projecting cells consistently clustered in dorsal and ventral aspects of RE in rostromedial levels (right). Blue squares represent regions in which clusters were found and further analyzed. Scale bar = 300μm. **C**: Percentage of retrogradely labeled RE to mPFC, REto vCA1 and dual mPFC-HC projecting neurons in RE dorsal and ventral regions of interest. All percentages are over total number of DAPI cells. **D-E:** Merged captures of immunofluorescent RE dual labeled cells **(D_i_-E_ii_)** and RE DAB CR^+^ **(D_i_, D_ii_)** or DAB CB^+^ **(E_i_,E_ii_)** cell bodies in dorsal and ventral RE. For process see Fig. S1. No dual labeling between dual mPFC-HC projecting cells and CR^+^ or CB^+^ cells in RE was observed. The relative distance between the center of RE dual labeled cells (yellow) and CR +DAB cells (marked with white ‘+’ signs) and/or CB+ DAB cells (marked with cyan ‘+’ signs) within a 100μm radius was measured using FIJI Image J. Scale bar = 50μm. **F-G:** Frequency distribution of DAB CR^+^ and CB^+^ cell counts in RE dorsal and ventral **(F_i_-G_ii_)** regions relative to radius distance (5μm bins) from RE dual (yellow) labeled cells. The number of DAB CR^+^ and DAB CB^+^ cells from RE dual labeled cells increased as a function of distance as shown by the linear regression line (black lines over histograms). Asterisks indicate significance: ** *p* < .01; *\*\*\*p* < .001 Abbreviations: rAAV, retrograde adeno-associated virus; CA1, CA1 subfield of the hippocampus; CAG, chicken beta-Actin promoter; CB, calbindin; CR, calretinin; DAB, 3,3’-Diaminobenzidine; DAPI, 4’,6-Diamidino-2-phenylindole dihydrochloride; GFP, green fluorescent protein; HC, hippocampus; IL, infralimbic cortex; mPFC, medial prefrontal cortex; PL, prelimbic cortex; tdTomato, red fluorescent protein; vCA1, ventral CA1.

## DISCUSSION

### Summary of Main Findings

The present study examined the calcium binding protein (PV, CR and CB) organization of the midline thalamus focusing on PVT, PT, RH and RE using a dual-labeling immunofluorescence approach. We further targeted specific dual mPFC-RE projecting cells in RE using a dual retrograde AAV tracing technique because these cells are theoretically critical to synchronous mPFC and HC activity (Dolleman-van der Weel et al., 2019; Hoover & Vertes, 2012; Varela et al., 2014). First, we did not find any PV^+^ cells in any of the nuclei of the midline thalamus, consistent with previous reports (Celio, 1990; Arai et al., 1994; Bokor et al., 2002). However, we did find an abundance of PV^+^ fibers in midline thalamus which are known to be inhibitory afferents from TRN (Albéri et al., 2013; Arai et al., 1994; McKenna & Vertes, 2004). Next, we demonstrated that CR^+^ and CB^+^ labeling organized the midline thalamus into distinct cell-type dominant zones. Notably, the dorsal and ventral CR^+^ and CB^+^ patterns mirrored each other suggesting a common developmental trajectory of CR^+^ and CB^+^ cell densities throughout the midline thalamus (Frassoni et al., 1998) in a pattern that simply flipped around the curvature of the rostral thalamus and 3V (e.g., vertically flipping Fig. 2C aligns the cell distributions of RE exceptionally well with those in Fig. 2B). That is, the dorsal midline thalamus (PVT and PT) contained a high density of CR^+^ cells and fibers that resembled a “T” or “Y” bordering the walls of the dorsal 3V and populating the midline. Whereas, CB^+^ cells were populated ventrolaterally and dense in PT. The ventral midline thalamus (RH and RE) contained a high density of CR^+^ cells resembling an inverted “T” or “Y” bordering the walls of the ventral 3V and occupied the midline including the center of rostral RH. CB^+^ cells in ventral midline thalamus were situated dorsolaterally and in lateral and caudal RH. Throughout the midline thalamus, dual-labeled CR^+^/CB^+^ cells were contained in partially overlapping single-labeled CR^+^ and CB^+^ zones. We detailed subregional variations on these patterns throughout the results, noting RE had the most complexity (summarized in Fig. 7). While we found a consistently opposing pattern of CR and CB cell density in PVT and RE across the rostro-caudal axis of the thalamus, with CR^+^ cell density higher in PVT and CB^+^ cell density was higher in RE, these differences may be, in part, due to the lack of an atlas separation for the CB^+^ cell population in the dorsolateral areas of RE in the way that PT is separated from PVT. Lastly, we showed that dual mPFC-HC projecting cells labeled with neither CR or CB, but were surrounded by rings of cells expressing both calcium binding proteins (composed of CR^+^, CB^+^, and CR^+^/CB^+^ cells). These dual mPFC-HC projecting center-ring organizations are potentially important microcircuits for midline thalamic integration and mPFC-HC synchronization.

**Figure 7.**
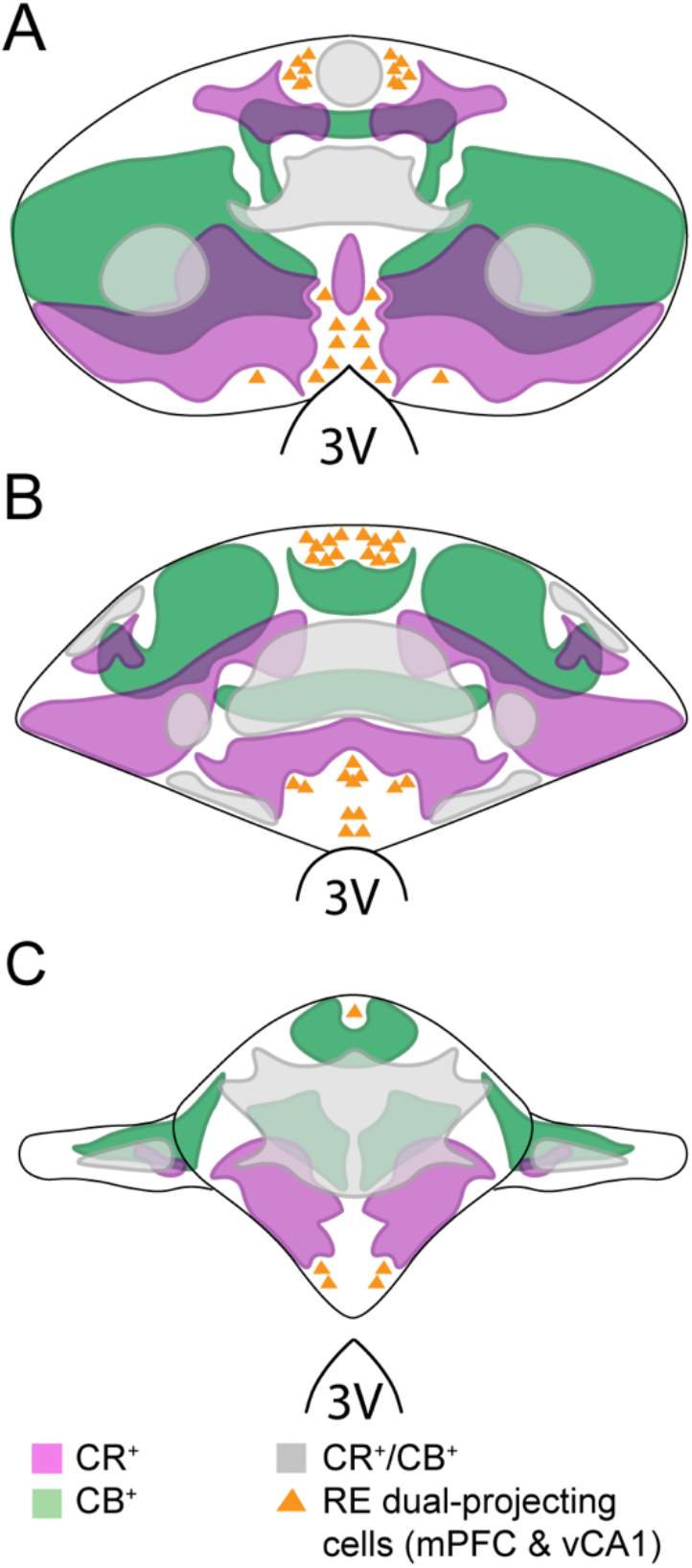
Topographical localization of CR and CB cell zones relative to dual mPFC-vHC projecting cells within RE. Schematic representation of the distribution of CR^+^ and CB^+^ cells in RE across **(A)** rostral, **(B)** medial, and **(C)** caudal levels. Colored areas represent zones where CR^+^ (light magenta) and CB^+^ (green) cells are concentrated. Intense magenta color represent regions in which CB and CR are in the same region, but do not overlap. Gray regions depict zones with dual labeled CR^+^/CB^+^ cells are predominantly seen. Yellow triangles represent dual mPFC-HC projecting cells. Overlay shown adapted from Swanson (2018). Abbreviations: CB, calbindin; CR, calretinin; mPFC, medial prefrontal cortex; RE, nucleus reuniens; vHC, ventral hippocampus; vCA1, ventral CA1; 3V, third ventricle.

### Calcium Binding Protein Distributions in Midline Thalamus

Overall, our results are in strong agreement with the outcomes of other studies that looked at the distribution of the calcium binding proteins in the rat thalamus that indicated that the midline thalamus is particularly rich/dense with CR^+^ and CB^+^ cells (Arai et al., 1994; Bokor et al., 2002; Winsky et al., 1992). However, we detail a few important differences with respect to the findings described in Arai et al. (1994), who productively used a single-labeling chromagen approach, on the distribution of the CR^+^ and CB^+^ cells in PVT, RH, and RE (our PT findings were nearly identical). First, Arai et al. (1994) indicated PVT was mainly a CR-containing structure while we found both CB^+^ and dual-labeled CR^+^/CB^+^ cells common throughout the region. PVT CR-labeling intensity and cells sizes were notably greater than for CB in PVT. Second, Arai et al., (1994) showed RH was mostly a CR-containing structure, but our data demonstrates that this is only true at the most rostromedial levels of RH. In stark contrast, we found that CB^+^ cells were almost exclusively populated at lateral and caudal levels of RH. Third, with respect to RE, Arai et al. (1994) described this nucleus as containing similar densities of CR and CB, while we demonstrate that RE contained more CB^+^ than CR^+^ cells and exhibits several distinct CR^+^, CB^+^ and CR^+^/CB^+^ cell zones (see Fig. 7 for details). The differences we found are likely due to the fact that we used a dual-labeling immunofluorescence protocol that was (1) more able to pick up on detailed distributions of CR^+^ and CB^+^ labeling in direct relationship to each other, and (2) confirm the identity of dual-labeled CR^+^/CB^+^ cells with a confocal z-stack analysis.

### Functional Implication of CR^+^ and CB^+^ Topographies in Midline Thalamus

It well known that calcium binding proteins such as CR^+^ and CB^+^ are crucially involved in neuronal functions. While CR and CB have been chiefly classified as slow buffers, recent work suggest they can also act as calcium sensors (Nelson & Chazin, 1998; Schwaller, 2014) Generally, calcium binding proteins have been used as an important tool in differentiating various cell types in the brain (Andressen et al., 1993; Arai et al., 1994; DeFelipe, 1997; Gulyás et al., 1996; Jones & Hendry, 1989). In fact, an early classification of the thalamus separated cells based on their calcium binding protein expression (Jones, 1998; Jones, 2001). Of these, CB^+^ cells in midline thalamus were classified as ‘matrix’ cells because of their dedicated projections to multimodal sensory regions in superficial layers of the cortex; and PV^+^ cells were considered ‘core’ cells for their projections to areas involved in the processing of sensory or motor information in middle cortical layers. While we did not find any PV^+^ cells in midline thalamus, CB^+^ cells were abundant especially in PT, RH and dorsolateral RE. The midline thalamus might be considered composed entirely of matrix cells involved in the processing of information from multiple brain regions in much the same way promoting cortico-thalamo-cortico synchronization, although the value of this distinction is not clear for further understanding subregional microcircuits in the midline thalamus without further investigation (Dolleman-van der Weel et al., 2019). While a role of CR is less known at this level of analysis, a population of CR^+^ RE cells that project to entorhinal cortex has been described (Wouterlood et al., 2007). We assume this is reflected in our observation of very dense and prominent CR^+^ fibers that extended throughout the shallow layers of entorhinal cortex but did not extend into perirhinal cortex, or into other cortices, which appeared to have more CB^+^ labeling. Given these entorhinal projections, CR^+^ cells in midline thalamus also seem to target multimodal sensory regions of cortex like CB^+^ cells. This suggest that the existence of separate and unique CR and CB zones in midline thalamus represent separate thalamo-cortical circuits.

Distinct CR^+^, CB^+^ and CR^+^/CB^+^ topographical zones in midline thalamus are likely associated with distinct rhythmicity like delta (e.g., Roy et al., 2017; Ferraris et al., 2018; Todorova and Zugaro, 2019; Schultheiss et al., 2020), theta (Hallock et al., 2016; Hasselmo et al., 2002; Jankowski et al., 2014; Lara-Vásquez et al., 2016; Roy et al., 2017; Vertes et al., 2004), or with sharp-wave ripples (Jadhav et al., 2012; Jadhav et al., 2016). Good support for this notion stems from the demonstration that CR^+^ cells in midline thalamus exhibit distinct *in vivo* electrophysiological profiles (Lara-Vásquez et al., 2016). Specifically, Lara-Vasquez et al. (2016) found that CR^+^ cells were more prone to bursting, not recruited in HC theta states, and inhibited by sharp-wave ripples. They also found that CR^-^ neurons were less prone to bursting and had no apparent relationship to theta (regardless of labeling for CB). These cells were sampled from across the dorsal-ventral extent of the midline thalamus signifying a primal role for calcium binding protein status. Combined with the present results, this suggests special considerations should be given when recording from midline thalamic neurons as to their calcium binding protein identity. For example, activity recorded from the CR-rich zones in RE located along the midline or near the third ventricle will likely differ significantly from RE activity recorded from the CB-rich dorsolateral areas, particularly with respect to their rhythmic and bursting profiles.

### Dual mPFC-HC projecting RE cells are distinct from CR^+^ and CB^+^ cell populations

Using a retrograde AAV viral approach we observed distinct and stereotyped clusters of RE cells with monosynaptic projections both to mPFC (prelimbic and infralimbic area) and ventral CA1. This finding is in agreement with two other studies that used more traditional tracing techniques (Hoover & Vertes, 2012; Varela et al., 2014). Specifically, Hoover and Vertes (2012) found several clusters of dual labeled cells in RE following injections of the retrograde tracers fluorogold and fluororuby in mPFC (PL/IL) and dorsal CA1, and in vCA1 and subiculum. Hoover and Vertes (2012) suggested that the functional roles of these cells were to support limbic subcortical and cortical interactions and/or the convergence and integration among other limbic-related structures. Varela et al. (2014) also demonstrated the existence of dual mPFC-HC projecting cells in RE following injections of cholera toxin (CTB) tracers. Varela et al. (2014) speculated these cells may be critical for the synchronization of target regions during exploration, transfer of mnemonic information between mPFC-HC, and modulation of certain phases of cortical spindle oscillations. It’s notable that we found that the dual projecting cells lacked CR or CB, but were surrounded by a ring of CR^+^, CB^+^ and CR^+^/CB^+^ cells. That is, we observed CB^+^, CR^+^ and CR^+^/CB^+^ cells formed well-defined circular clusters in regions that were sparse or seemly absent of CR^+^ or CB^+^ cells (within a 100μm radius). Reliably, CR and CB cell densities were scant near dual mPFC-HC projecting cells, but cell densities rapidly increased with distance. Speculatively, this organization may provide a microcircuit means for integrating input activity from several different cell types and then synchronizing outputs to mPFC and CA1 for triggering coordinated rhythmic modes (e.g., delta or theta). Interrogating these cells further will require sophisticated functional approaches such as optogenetics delivered with combinatorial retrograde and Cre-dependent viral constructs.

### Conclusion

Most often, neurons in midline thalamus are assumed to be a relatively homogeneous group of excitatory (glutamatergic) projection neurons (Bokor et al., 2002), but here we detailed several distinct zones in the midline thalamus based on the expression of CR and CB, or lack thereof in dual mPFC-HC projecting cells. While experiments have productively targeted entire regions of the midline thalamus, most commonly RH/RE with inactivations or lesions (Cholvin et al., 2013; Hallock et al., 2013; Hembrook et al., 2012; Layfield et al., 2015; Loureiro et al., 2012), we suggest it may be more productive for future experiments to separately target CR^+^, CB^+^, and dual mPFC-HC projecting cells to better understand the computational and/or rhythmic contributions of the midline thalamus to the mPFC-HC system.

## Acknowledgements

The authors would like to thank Amanda Pacheco-Spiewak and Nashya Linares for their assistance in cell counting; Maanasa Jayachandran and Maximilian Schlecht for experimental assistance; and Jennifer Dziedzic and Tomás Guilarte for help with microscopy. This work was supported, in part, by NIH grant MH113626 to TAA, and funds from the FIU CASE Distinguished Postdoctoral Program to TDV.

## REFERENCES

Ahn, J. H., Hong, S., Park, J. H., Kim, I. H., Cho, J. H., Lee, T.-K., Lee, J.-C., Chen, B. H., Shin, B.-N., Bae, E. J., Jeon, Y. H., Kim, Y.-M., Won, M.-H., & Choi, S. Y. (2017). Immunoreactivities of calbindin-D28k, calretinin and parvalbumin in the somatosensory cortex of rodents during normal aging. Molecular Medicine Reports, 16(5), 7191–7198. https://doi.org/10.3892/mmr.2017.7573

Aika, Y., Ren, J. Q., Kosaka, K., & Kosaka, T. (1994). Quantitative analysis of GABA-like-immunoreactive and parvalbumin-containing neurons in the CA1 region of the rat hippocampus using a stereological method, the disector. Experimental Brain Research, 99(2). https://doi.org/10.1007/BF00239593

Al-Mashhadi, S., Simpson, J. E., Heath, P. R., Dickman, M., Forster, G., Matthews, F. E., Brayne, C., Ince, P. G., Wharton, S. B., & Medical Research Council Cognitive Function and Ageing Study. (2015). Oxidative glial cell damage associated with white matter lession in the aging human brain: Oxidative damage in white matter lesions. Brain Pathology, 25(5), 565–574. https://doi.org/10.1111/bpa.12216

Albéri, L., Lintas, A., Kretz, R., Schwaller, B., & Villa, A. E. P. (2013). The calcium-binding protein parvalbumin modulates the firing 1 properties of the reticular thalamic nucleus bursting neurons. Journal of Neurophysiology, 109(11), 2827–2841. https://doi.org/10.1152/jn.00375.2012

Andressen, C., Blümcke, I., & Celio, M. R. (1993). Calcium-binding proteins: Selective markers of nerve cells. Cell and Tissue Research, 271(2), 181–208. https://doi.org/10.1007/BF00318606

Anaconda Software Distribution. (2016). Anaconda (Version 2-2.4.0). [Computer software]. Retrieved from: https://anaconda.com

Arai, R., Jacobowitz, D. M., & Deura, S. (1994). Distribution of calretinin, calbindin-D28k, and parvalbumin in the rat thalamus. Brain Research Bulletin, 33(5), 595–614. https://doi.org/10.1016/0361-9230(94)90086-8

Barker, G. R. I., & Warburton, E. C. (2018). A critical role for the nucleus reuniens in long-rerm, but not short-term associative recognition memory formation. The Journal of Neuroscience, 38(13), 3208–3217. https://doi.org/10.1523/JNEUROSCI.1802-17.2017

Bokor, H., Csáki, Á., Kocsis, K., & Kiss, J. (2002). Cellular architecture of the nucleus reuniens thalami and its putative aspartatergic/glutamatergic projection to the hippocampus and medial septum in the rat: Reuniens connection to hippocampus and septum. European Journal of Neuroscience, 16(7), 1227–1239. https://doi.org/10.1046/j.1460-9568.2002.02189.x

Braak, H., & Braak, E. (1991). Alzheimer’s disease affects limbic nuclei of the thalamus. Acta Neuropathologica, 81(3), 261–268. https://doi.org/10.1007/BF00305867

Burwell, R. D. (2000). The parahippocampal region: Corticocortical connectivity. Annals of the New York Academy of Sciences, 911, 25–42. https://doi.org/10.1111/j.1749-6632.2000.tb06717.x

Cassel, J.-C., de Vasconcelos, A., Loureiro, M., Cholvin, T., Dalrymple-Alford, J. C., & Vertes, R. P. (2013). The reuniens and rhomboid nuclei: Neuroanatomy, electrophysiological characteristics and behavioral implications. Progress in Neurobiology, 111, 34–52. https://doi.org/10.1016/j.pneurobio.2013.08.006

Celio, M. R. (1990). Calbindin D-28k and parvalbumin in the rat nervous system. Neuroscience, 35(2), 375–475. https://doi.org/10.1016/0306-4522(90)90091-H

Cenquizca, L. A., & Swanson, L. W. (2007). Spatial organization of direct hippocampal field CA1 axonal projections to the rest of the cerebral cortex. Brain Research Reviews, 56(1), 1–26. https://doi.org/10.1016/j.brainresrev.2007.05.002

Choi, E. A., & McNally, G. P. (2017). Paraventricular thalamus balances danger and reward. The Journal of Neuroscience, 37(11), 3018–3029. https://doi.org/10.1523/JNEUROSCI.3320-16.2017

Cholvin, T., Loureiro, M., Cassel, R., Cosquer, B., Geiger, K., De Sa Nogueira, D., Raingard, H., Robelin, L., Kelche, C., de Vasconcelos, A., & Cassel, J.-C. (2013). The Ventral Midline Thalamus Contributes to Strategy Shifting in a Memory Task Requiring Both Prefrontal Cortical and Hippocampal Functions. Journal of Neuroscience, 33(20), 8772–8783. https://doi.org/10.1523/JNEUROSCI.0771-13.2013

Churchwell, J. C., & Kesner, R. P. (2011). Hippocampal-prefrontal dynamics in spatial working memory: Interactions and independent parallel processing. Behavioural Brain Research, 225(2), 389–395. https://doi.org/10.1016/j.bbr.2011.07.045

Condé, F., Lund, J. S., Jacobowitz, D. M., Baimbridge, K. G., & Lewis, D. A. (1994). Local circuit neurons immunoreactive for calretinin, calbindin D-28k or parvalbumin in monkey prefronatal cortex: Distribution and morphology: CALCIUM-BINDING PROTEINS IN MONKEY PREFRONTAL CORTEX. Journal of Comparative Neurology, 341(1), 95–116. https://doi.org/10.1002/cne.903410109

Csillik, B., Mihály, A., Krisztin-Péva, B., Chadaide, Z., Samsam, M., Knyihár-Csillik, E., & Fenyo, R. (2005). GABAergic parvalbumin-immunoreactive large calyciform presynaptic complexes in the reticular nucleus of the rat thalamus. Journal of Chemical Neuroanatomy, 30(1), 17–26. https://doi.org/10.1016/j.jchemneu.2005.03.010

DeFelipe, J. (1997). Types of neurons, synaptic connections and chemical characteristics of cells immunoreactive for calbindin-D28K, parvalbumin and calretinin in the neocortex. Journal of Chemical Neuroanatomy, 14(1), 1–19. https://doi.org/10.1016/S0891-0618(97)10013-8

del Río, M. R., & DeFelipe, J. (1996). Colocalization of calbindin D-28k, calretinin, and GABA immunoreactivities in neurons of the human temporal cortex. Journal of Comparative Neurology, 369(3), 472–482. https://doi.org/10.1002/(SICI)1096-9861(19960603)369:3<472::AID-CNE11>3.0.CO;2-K

Dolleman-van der Weel, M. J., Griffin, A. L., Ito, H. T., Shapiro, M. L., Witter, M. P., Vertes, R. P., & Allen, T. A. (2019). The nucleus reuniens of the thalamus sits at the nexus of a hippocampus and medial prefrontal cortex circuit enabling memory and behavior. Learning & Memory, 26(7), 191–205. https://doi.org/10.1101/lm.048389.118

Dolleman-van der Weel, M., & Witter, M. P. (1996). Projections from the nucleus reuniens thalami to the entorhinal cortex, hippocampal field CA1, and the subiculum in the rat arise from different populations of neurons. Journal of Comparative Neurology, 364(4), 637–650. https://doi.org/10.1002/(sici)1096-9861(19960122)364:4<637::aid-cne3>3.0.co;2-4

Eichenbaum, H. (2017). Prefrontal–hippocampal interactions in episodic memory. Nature Reviews Neuroscience, 18(9), 547–558. https://doi.org/10.1038/nrn.2017.74

Ferino, F., Thierry, A. M., & Glowinski, J. (1987). Anatomical and electrophysiological evidence for a direct projection from ammon’s horn to the medial prefrontal cortex in the rat. Experimental Brain Research, 65(2). https://doi.org/10.1007/BF00236315

Ferraris, M., Ghestem, A., Vicente, A. F., Nallet-Khosrofian, L., Bernard, C., & Quilichini, P. P. (2018). The nucleus reuniens controls long-range hippocampo–prefrontal gamma synchronization during slow oscillations. The Journal of Neuroscience, 38(12), 3026–3038. https://doi.org/10.1523/JNEUROSCI.3058-17.2018

Fonseca, M., & Soriano, E. (1995). Calretinin-immunoreactive neurons in the normal human temporal cortex and in Alzheimer’s disease. Brain Research, 691(1–2), 83–91. https://doi.org/10.1016/0006-8993(95)00622-W

Frassoni, C., Arcelli, P., Selvaggio, M., & Spreafico, R. (1998). Calretinin immunoreactivity in the developing thalamus of the rat: A marker of early generated thalamic cells. Neuroscience, 83(4), 1203–1214. https://doi.org/10.1016/S0306-4522(97)00443-0

Fuchs, E. C., Zivkovic, A. R., Cunningham, M. O., Middleton, S., LeBeau, F. E. N., Bannerman, D. M., Rozov, A., Whittington, M. A., Traub, R. D., Rawlins, J. N. P., & Monyer, H. (2007). Recruitment of Parvalbumin-Positive Interneurons Determines Hippocampal Function and Associated Behavior. Neuron, 53(4), 591–604. https://doi.org/10.1016/j.neuron.2007.01.031

Furtak, S. C., Wei, S.-M., Agster, K. L., & Burwell, R. D. (2007). Functional neuroanatomy of the parahippocampal region in the rat: The perirhinal and postrhinal cortices. Hippocampus, 17(9), 709–722. https://doi.org/10.1002/hipo.20314

Fuster, J. M. (1995). Cognitive Support of Behavior. Plasticity in the central nervous system: learning and memory, 149.

Gelinas, J. N., Khodagholy, D., Thesen, T., Devinsky, O., & Buzsáki, G. (2016). Interictal epileptiform discharges induce hippocampal–cortical coupling in temporal lobe epilepsy. Nature Medicine, 22(6), 641–648. https://doi.org/10.1038/nm.4084

Gulyás, A. I., Hájos, N., & Freund, T. F. (1996). Interneurons Containing Calretinin Are Specialized to Control Other Interneurons in the Rat Hippocampus. The Journal of Neuroscience, 16(10), 3397–3411. https://doi.org/10.1523/JNEUROSCI.16-10-03397.1996

Hallock, H L, Wang, A., & Griffin, A. L. (2016). Ventral Midline Thalamus Is Critical for Hippocampal-Prefrontal Synchrony and Spatial Working Memory. Journal of Neuroscience, 36(32), 8372–8389. https://doi.org/10.1523/JNEUROSCI.0991-16.2016

Hallock, Henry L, Wang, A., Shaw, C. L., & Griffin, A. L. (2013). Transient inactivation of the thalamic nucleus reuniens and rhomboid nucleus produces deficits of a working-memory dependent tactile-visual conditional discrimination task. Behavioral Neuroscience, 127(6), 860–866. https://doi.org/10.1037/a0034653

Hasselmo, M. E., Bodelón, C., & Wyble, B. P. (2002). A proposed function for hippocampal theta rhythm: Separate phases of encoding and retrieval enhance reversal of prior learning. Neural Computation, 14(4), 793–817. https://doi.org/10.1162/089976602317318965

Hauer, B. E., Pagliardini, S., & Dickson, C. T. (2019). The Reuniens Nucleus of the Thalamus Has an Essential Role in Coordinating Slow-Wave Activity between Neocortex and Hippocampus. Eneuro, 6(5), ENEURO.0365--19.2019. https://doi.org/10.1523/ENEURO.0365-19.2019

Hembrook, J. R., Onos, K. D., & Mair, R. G. (2012). Inactivation of ventral midline thalamus produces selective spatial delayed conditional discrimination impairment in the rat. Hippocampus, 22(4), 853–860. https://doi.org/10.1002/hipo.20945

Hof, P. R., Glezer, I. I., Condé, F., Flagg, R. A., Rubin, M. B., Nimchinsky, E. A., & Vogt Weisenhorn, D. M. (1999). Cellular distribution of the calcium-binding proteins parvalbumin, calbindin, and calretinin in the neocortex of mammals: phylogenetic and developmental patterns. Journal of Chemical Neuroanatomy, 16(2), 77–116. https://doi.org/10.1016/S0891-0618(98)00065-9

Hoover, W. B., & Vertes, R. P. (2012). Collateral projections from nucleus reuniens of thalamus to hippocampus and medial prefrontal cortex in the rat: A single and double retrograde fluorescent labeling study. Brain Structure and Function, 217(2), 191–209. https://doi.org/10.1007/s00429-011-0345-6

Hsu, D. T., Kirouac, G. J., Zubieta, J.-K., & Bhatnagar, S. (2014). Contributions of the paraventricular thalamic nucleus in the regulation of stress, motivation, and mood. Frontiers in Behavioral Neuroscience, 8. https://doi.org/10.3389/fnbeh.2014.00073

Hunsaker, M. R., & Kesner, R. P. (2018). Unfolding the cognitive map: The role of hippocampal and extra-hippocampal substrates based on a systems analysis of spatial processing. Neurobiology of Learning and Memory, 147, 90–119. https://doi.org/10.1016/j.nlm.2017.11.012

Ito, H. T., Zhang, S.-J., Witter, M. P., Moser, E. I., & Moser, M.-B. (2015). A prefrontal–thalamo–hippocampal circuit for goal-directed spatial navigation. Nature, 522(7554), 50–55. https://doi.org/10.1038/nature14396

Jadhav, S. P., Kemere, C., German, P. W., & Frank, L. M. (2012). Awake hippocampal sharp-wave ripples support spatial memory. Science, 336(6087), 1454–1458. https://doi.org/10.1126/science.1217230

Jadhav, S. P. P., Rothschild, G., Roumis, D. K. K., & Frank, L. M. M. (2016). Coordinated Excitation and Inhibition of Prefrontal Ensembles during Awake Hippocampal Sharp-Wave Ripple Events. Neuron, 90(1), 113–127. https://doi.org/10.1016/j.neuron.2016.02.010

Jankowski, M. M., Islam, M. N., Wright, N. F., Vann, S. D., Erichsen, J. T., Aggleton, J. P., & O’Mara, S. M. (2014). Nucleus reuniens of the thalamus contains head direction cells. ELife, 3(July 2014), 1–10. https://doi.org/10.7554/eLife.03075

Jayachandran, M., Linley, S. B., Schlecht, M., Mahler, S. V, Vertes, R. P., & Allen, T. A. (2019). Prefrontal Pathways Provide Top-Down Control of Memory for Sequences of Events. Cell Reports, 28(3), 640--654.e6. https://doi.org/10.1016/j.celrep.2019.06.053

Jin, J., & Maren, S. (2015). Prefrontal-hippocampal interactions in memory and emotion. Frontiers in Systems Neuroscience, 9. https://doi.org/10.3389/fnsys.2015.00170

Jones, E G. (1998). Viewpoint: the core and matrix of thalamic organization. Neuroscience, 85(2), 331–345. https://doi.org/10.1016/S0306-4522(97)00581-2

Jones, E G, & Hendry, S. H. C. (1989). Differential calcium binding protein immunoreactivity distinguishes classes of relay neurons in monkey thalamic nuclei. European Journal of Neuroscience, 1(3), 222–246. https://doi.org/10.1111/j.1460-9568.1989.tb00791.x

Jones, Edward G. (2001). The thalamic matrix and thalamocortical synchrony. Trends in Neurosciences, 24(10), 595–601. https://doi.org/10.1016/S0166-2236(00)01922-6

Kawano, J. (2001). Suprachiasmatic nucleus projections to the paraventricular thalamic nucleus of the rat. Thalamus & Related Systems, 1(3), 197–202. https://doi.org/10.1016/S1472-9288(01)00019-X

Kerr, K. M., Agster, K. L., Furtak, S. C., & Burwell, R. D. (2007). Functional neuroanatomy of the parahippocampal region: The lateral and medial entorhinal areas. Hippocampus, 17(9), 697–708. https://doi.org/10.1002/hipo.20315

Kirichenko, E. Y., Matsionis, A. E., Povilaitite, P. E., Akimenko, M. A., & Logvinov, A. K. (2017). Characteristics of the Structural Organization of the Ventral Posteromedial and Posterolateral Nuclei and the Reticular Nucleus of the Thalamus in Rats (an immunohistochemical study). Neuroscience and Behavioral Physiology, 47(6), 621–626. https://doi.org/10.1007/s11055-017-0444-9

Kosaka, T., Wu, J.-Y., & Benoit, R. (1988). GABAergic neurons containing somatostatin-like immunoreactivity in the rat hippocampus and dentate gyrus. Experimental Brain Research, 71(2). https://doi.org/10.1007/BF00247498

Lara-Vásquez, A., Espinosa, N., Durán, E., Stockle, M., & Fuentealba, P. (2016). Midline thalamic neurons are differentially engaged during hippocampus network oscillations. Scientific Reports, 6(1), 29807. https://doi.org/10.1038/srep29807

Layfield, D. M., Patel, M., Hallock, H., & Griffin, A. L. (2015). Inactivation of the nucleus reuniens/rhomboid causes a delay-dependent impairment of spatial working memory. Neurobiology of Learning and Memory, 125, 163–167. https://doi.org/10.1016/j.nlm.2015.09.007

Li, S., & Kirouac, G. J. (2008). Projections from the paraventricular nucleus of the thalamus to the forebrain, with special emphasis on the extended amygdala. The Journal of Comparative Neurology, 506(2), 263–287. https://doi.org/10.1002/cne.21502

Lisman, J. E., Pi, H. J., Zhang, Y., & Otmakhova, N. A. (2010). A Thalamo-Hippocampal-Ventral Tegmental Area Loop May Produce the Positive Feedback that Underlies the Psychotic Break in Schizophrenia. Biological Psychiatry, 68(1), 17–24. https://doi.org/10.1016/j.biopsych.2010.04.007

Loureiro, M., Cholvin, T., Lopez, J., Merienne, N., Latreche, A., Cosquer, B., Geiger, K., Kelche, C., Cassel, J.-C., & de Vasconcelos, A. (2012). The Ventral Midline Thalamus (Reuniens and Rhomboid Nuclei) Contributes to the Persistence of Spatial Memory in Rats. Journal of Neuroscience, 32(29), 9947–9959. https://doi.org/10.1523/JNEUROSCI.0410-12.2012

Majercikova, Z., Weering, H. van, Scsukova, S., Mikkelsen, J. D., & Kiss, A. (2012). A new approach of light microscopic immunohistochemical triple-staining: combination of Fos labeling with diaminobenzidine-nickel and neuropeptides labeled with Alexa488 and Alexa555 fluorescent dyes. Endocrine Regulations, 46(04), 217–223. https://doi.org/10.4149/endo_2012_04_217

Matzeu, A., Zamora-Martinez, E. R., & Martin-Fardon, R. (2014). The paraventricular nucleus of the thalamus is recruited by both natural rewards and drugs of abuse: recent evidence of a pivotal role for orexin/hypocretin signaling in this thalamic nucleus in drug-seeking behavior. Frontiers in Behavioral Neuroscience, 8. https://doi.org/10.3389/fnbeh.2014.00117

McGaugh, J. L., Bermudez-Rattoni, F., & Prado-Alcala, R. A. (2019). Plasticity in the central nervous system: Learning and memory. In Plasticity in the Central Nervous System: Learning and Memory. https://doi.org/10.4324/9781315789279

McKenna, J. T., & Vertes, R. P. (2004). Afferent projections to nucleus reuniens of the thalamus. Journal of Comparative Neurology, 480(2), 115–142. https://doi.org/10.1002/cne.20342

McQuin, C., Goodman, A., Chernyshev, V., Kamentsky, L., Cimini, B. A., Karhohs, K. W., Doan, M., Ding, L., Rafelski, S. M., Thirstrup, D., Wiegraebe, W., Singh, S., Becker, T., Caicedo, J. C., & Carpenter, A. E. (2018). CellProfiler 3.0: Next-generation image processing for biology. PLOS Biology, 16(7), e2005970. https://doi.org/10.1371/journal.pbio.2005970

Miettinen, R., Gulyás, A. I., Baimbridge, K. G., Jacobowitz, D. M., & Freund, T. F. (1992). Calretinin is present in non-pyramidal cells of the rat hippocampus—II. Co-existence with other calcium binding proteins and gaba. Neuroscience, 48(1), 29–43. https://doi.org/10.1016/0306-4522(92)90335-Y

Moyer, J. R., Furtak, S. C., McGann, J. P., & Brown, T. H. (2011). Aging-related changes in calcium-binding proteins in rat perirhinal cortex. Neurobiology of Aging, 32(9), 1693–1706. https://doi.org/10.1016/j.neurobiolaging.2009.10.001

Nelson, A. J. D., Hindley, E. L., Haddon, J. E., Vann, S. D., & Aggleton, J. P. (2014). A novel role for the rat retrosplenial cortex in cognitive control. Learning & Memory, 21(2), 90–97. https://doi.org/10.1101/lm.032136.113

Nelson, M. R., & Chazin, W. J. (1998). Structures of EF-hand Ca2+-binding proteins: Diversity in the organization, packing and response to Ca2+ binding. BioMetals, 11(4), 297–318. https://doi.org/10.1023/A:1009253808876

Penzo, M. A., Robert, V., Tucciarone, J., De Bundel, D., Wang, M., Van Aelst, L., Darvas, M., Parada, L. F., Palmiter, R. D., He, M., Huang, Z. J., & Li, B. (2015). The paraventricular thalamus controls a central amygdala fear circuit. Nature, 519(7544), 455–459. https://doi.org/10.1038/nature13978

Preston, A. R., & Eichenbaum, H. (2013). Review Interplay of Hippocampus and Prefrontal Cortex in Memory. CURBIO, 23, R764--R773. https://doi.org/10.1016/j.cub.2013.05.041

Reynolds, G. P., Abdul-Monim, Z., Neill, J. C., & Zhang, Z.-J. (2004). Calcium binding protein markers of GABA deficits in schizophrenia — post mortem studies and animal models. Neurotoxicity Research, 6(1), 57–61. https://doi.org/10.1007/BF03033297

Rogers, J. H., & Résibois, A. (1992). Calretinin and calbindin-D28k in rat brain: Patterns of partial co-localization. Neuroscience, 51(4), 843–865. https://doi.org/10.1016/0306-4522(92)90525-7

Roy, A., Svensson, F. P., Mazeh, A., & Kocsis, B. (2017). Prefrontal-hippocampal coupling by theta rhythm and by 2–5 Hz oscillation in the delta band: The role of the nucleus reuniens of the thalamus. Brain Structure and Function, 222(6), 2819–2830. https://doi.org/10.1007/s00429-017-1374-6

Schultheiss, N. W., Schlecht, M., Jayachandran, M., Brooks, D. R., McGlothan, J. L., Guilarte, T. R., & Allen, T. A. (2020). Awake delta and theta-rhythmic hippocampal network modes during intermittent locomotor behaviors in the rat. Behavioral Neuroscience, undefined(undefined), undefined. https://doi.org/10.1037/bne0000409

Schwaller, B. (2014). Calretinin: from a “simple” Ca2+ buffer to a multifunctional protein implicated in many biological processes. Frontiers in Neuroanatomy, 8. https://doi.org/10.3389/fnana.2014.00003

Sherman, S. M. (2017). Functioning of Circuits Connecting Thalamus and Cortex. In R. Terjung (Ed.), Comprehensive Physiology (pp. 713–739). John Wiley & Sons, Inc. https://doi.org/10.1002/cphy.c160032

Schindelin, J., Arganda-Carreras, I., Frise, E., Kaynig, V., Longair, M., Pietzsch, T., Preibisch, S., Rueden, C., Saalfeld, S., Schmid, B., Tinevez, J.Y., White, D. J., Hartenstein, V., Eliceiri, K., Tomancak, P., & Cardona, A. (2012). Fiji: An open-source platform for biological-image analysis. Nature Methods, 9(7), 676–682. https://doi.org/10.1038/nmeth.2019

Skelin, I., Kilianski, S., & McNaughton, B. L. (2019). Hippocampal coupling with cortical and subcortical structures in the context of memory consolidation. Neurobiology of Learning and Memory, 160, 21–31. https://doi.org/10.1016/j.nlm.2018.04.004

Sloviter, R. S. (1989). Calcium-binding protein (calbindin-D28k) and parvalbumin immunocytochemistry: Localization in the rat hippocampus with specific reference to the selective vulnerability of hippocampal neurons to seizure activity. The Journal of Comparative Neurology, 280(2), 183–196. https://doi.org/10.1002/cne.902800203

Spellman, T., Rigotti, M., Ahmari, S. E., Fusi, S., Gogos, J. A., & Gordon, J. A. (2015). Hippocampal–prefrontal input supports spatial encoding in working memory. Nature, 522(7556), 309–314. https://doi.org/10.1038/nature14445

Su, H.-S., & Bentivoglio, M. (1990). Thalamic midline cell populations projecting to the nucleus accumbens, amygdala, and hippocampus in the rat. The Journal of Comparative Neurology, 297(4), 582–593. https://doi.org/10.1002/cne.902970410

Swanson, L. W. (2018). Brain maps 4.0-Structure of the rat brain: An open access atlas with global nervous system nomenclature ontology and flatmaps. Journal of Comparative Neurology, 526(6), 935–943. https://doi.org/10.1002/cne.24381

Todorova, R., & Zugaro, M. (2019). Isolated cortical computations during delta waves support memory consolidation. Science, 366(6463), 377–381. https://doi.org/10.1126/science.aay0616

Tollemar, V., Tudzarovski, N., Boberg, E., Törnqvist Andrén, A., Al-Adili, A., Le Blanc, K., Garming Legert, K., Bottai, M., Warfvinge, G., & Sugars, R. V. (2018). Quantitative chromogenic immunohistochemical image analysis in cellprofiler software: Quantitative chromogenic immunohistochemistry. Cytometry Part A, 93(10), 1051–1059. https://doi.org/10.1002/cyto.a.23575

Urbán, Z., Maglóczky, Z., & Freund, T. F. (2002). Calretinin-containing interneurons innervate both principal cells and interneurons in the CA1 region of the human hippocampus. Acta Biologica Hungarica, 53(1–2), 205–220. https://doi.org/10.1556/ABiol.53.2002.1-2.19

Van Brederode, J. F. M., Helliesen, M. K., & Hendrickson, A. E. (1991). Distribution of the calcium-binding proteins parvalbumin and calbindin-D28k in the sensorimotor cortex of the rat. Neuroscience, 44(1), 157–171. https://doi.org/10.1016/0306-4522(91)90258-P

Varela, C., Kumar, S., Yang, J. Y., & Wilson, M. A. (2014). Anatomical substrates for direct interactions between hippocampus, medial prefrontal cortex, and the thalamic nucleus reuniens. Brain Structure and Function, 219(3), 911–929. https://doi.org/10.1007/s00429-013-0543-5

Vertes, R. P., & Hoover, W. B. (2008). Projections of the paraventricular and paratenial nuclei of the dorsal midline thalamus in the rat. The Journal of Comparative Neurology, 508(2), 212–237. https://doi.org/10.1002/cne.21679

Vertes, R. P., Hoover, W. B., Do Valle, A. C., Sherman, A., & Rodriguez, J. J. (2006). Efferent projections of reuniens and rhomboid nuclei of the thalamus in the rat. Journal of Comparative Neurology, 499(5). https://doi.org/10.1002/cne.21135

Vertes, R. P., Hoover, W. B., Szigeti-Buck, K., & Leranth, C. (2007). Nucleus reuniens of the midline thalamus: Link between the medial prefrontal cortex and the hippocampus. Brain Research Bulletin, 71(6). https://doi.org/10.1016/j.brainresbull.2006.12.002

Vertes, R. P., Hoover, W. B., & Viana Di Prisco, G. (2004). Theta rhythm of the hippocampus: subcortical control and functional significance. In Behavioral and cognitive neuroscience reviews (Vol. 3, Issue 3, pp. 173–200). Behav Cogn Neurosci Rev. https://doi.org/10.1177/1534582304273594

Vertes, R. P., Linley, S. B., & Hoover, W. B. (2015). Limbic circuitry of the midline thalamus. In Neuroscience and Biobehavioral Reviews (Vol. 54). https://doi.org/10.1016/j.neubiorev.2015.01.014

Viena, T. D., Linley, S. B., & Vertes, R. P. (2018). Inactivation of nucleus reuniens impairs spatial working memory and behavioral flexibility in the rat. Hippocampus, 28(4). https://doi.org/10.1002/hipo.22831

Winsky, L., Montpied, P., Arai, R., Martin, B. M., & Jacobowitz, D. M. (1992). Calretinin distribution in the thalamus of the rat: Immunohistochemical and in situ hybridization histochemical analyses. Neuroscience, 50(1), 181–196. https://doi.org/10.1016/0306-4522(92)90391-E

Witter, M. P., Doan, T. P., Jacobsen, B., Nilssen, E. S., & Ohara, S. (2017). Architecture of the Entorhinal Cortex A Review of Entorhinal Anatomy in Rodents with Some Comparative Notes. Frontiers in Systems Neuroscience, 11, 46. https://doi.org/10.3389/fnsys.2017.00046

Wouterlood, F. G., Canto, C. B., Aliane, V., Boekel, A. J., Grosche, J., Härtig, W., Beliën, J. A. M., & Witter, M. P. (2007). Coexpression of vesicular glutamate transporters 1 and 2, glutamic acid decarboxylase and calretinin in rat entorhinal cortex. Brain Structure and Function, 212(3–4), 303–319. https://doi.org/10.1007/s00429-007-0163-z

Wouterlood, F. G., Grosche, J., & Härtig, W. (2001). Co-localization of calretinin and calbindin in distinct cells in the hippocampal formation of the rat. Brain Research, 922(2), 310–314. https://doi.org/10.1016/S0006-8993(01)03220-6

Xu, W., & Sudhof, T. C. (2013). A Neural Circuit for Memory Specificity and Generalization. Science, 339(6125), 1290–1295. https://doi.org/10.1126/science.1229534

Young, J. K., Wu, M., Manaye, K. F., Kc, P., Allard, J. S., Mack, S. O., & Haxhiu, M. A. (2005). Orexin stimulates breathing via medullary and spinal pathways. Journal of Applied Physiology, 98(4), 1387–1395. https://doi.org/10.1152/japplphysiol.00914.2004

Zimmer, D. B., Cornwall, E. H., Landar, A., & Song, W. (1995). The S100 protein family: History, function, and expression. Brain Research Bulletin, 37(4), 417–429. https://doi.org/10.1016/0361-9230(95)00040-2

