## Supplemental Materials for "Calretinin and calbindin architecture of the midline thalamus associated with prefrontal-hippocampal circuitry"

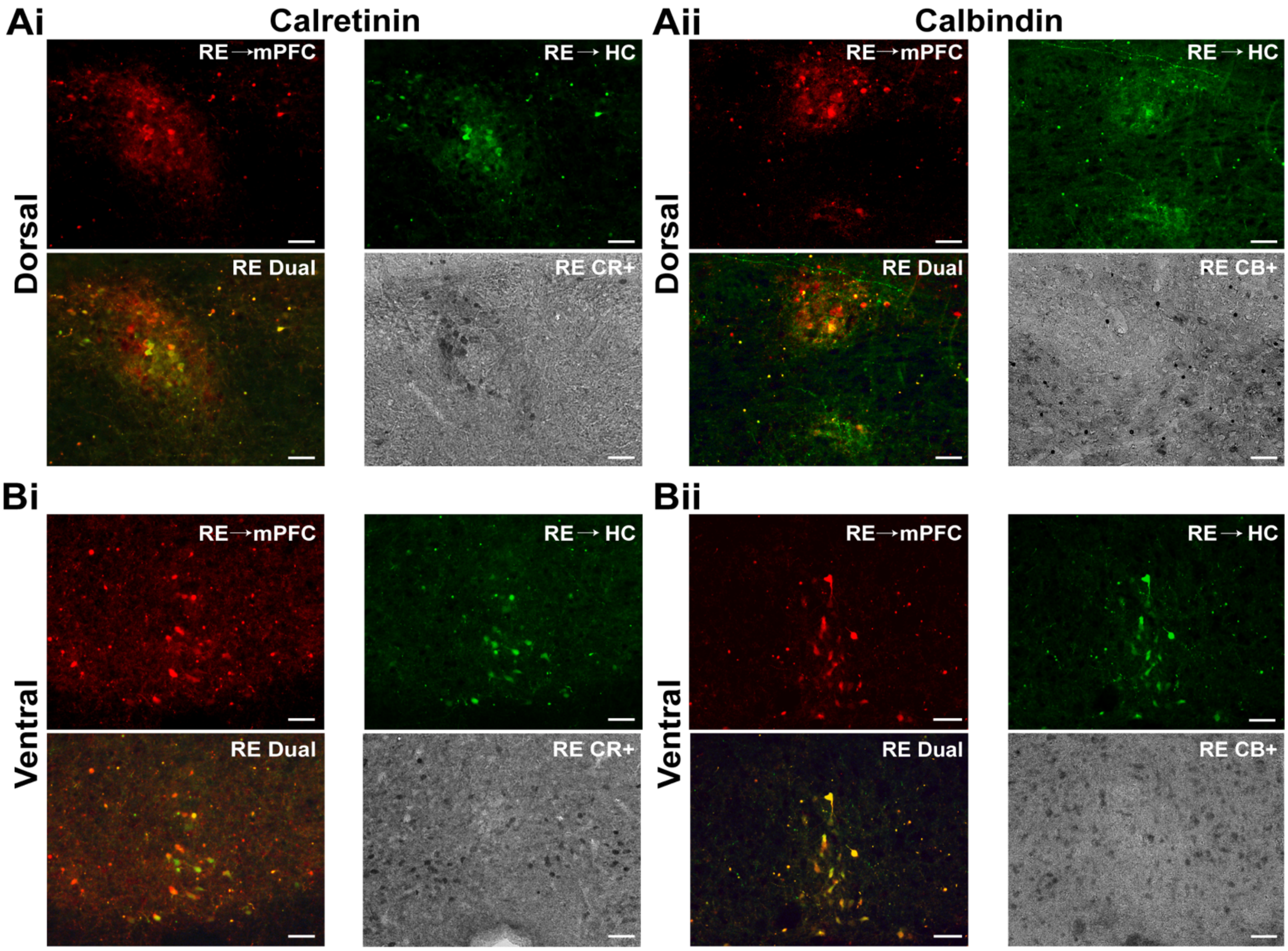


**Figure S1. Process of merging immunofluorescent and DAB images from a single site**

Representative images obtained in the same brain region across different channels using an epifluorescence/brightfield microscope. Using the epifluorescence mode, clusters of dual labeled RE neurons (yellow) were first identified in dorsal **($A_i$-$A_{ii}$)** and ventral **($B_i$-$B_{ii}$)**. Images of retrogradely labeled RE to mPFC cells were captured in the red channel, while RE to HC cells were captured in the green channel. Immediately after, using the brightfield mode, DAB RE $CR^+$ or RE $CB^+$ cells were captured. The red and green channel were merged, allowing the visualization of RE dual mPFC-HC projecting cells (yellow). Finally, the fluorescent merged image was transposed with its corresponding brightfield capture using Adobe Photoshop for the purpose of obtaining the final merged images shown in *Fig. 6*. Manual cell counts were done in each independent channel. Scale Bar = 50µm.

Abbreviations: CB, calbindin; CR, calretinin; DAB, 3,3'-Diaminobenzidine; mPFC, medial prefrontal cortex; RE, nucleus reuniens; HC, hippocampus

***Table S1. Mean soma size area of $CR^+$, $CB^+$ and $CR^+/CB^+$ neurons in paraventricular (PVT) and nucleus reuniens (RE)***

| Axis | Regional Subdivision | Calcium Binding Protein | PVT Mean (μm²) | PVT SD | RE Mean (μm²) | RE SD |
|---|---|---|---|---|---|---|
| Rostral | Dorsolateral | CR+ | 142.0 | 50.0 | 155.6 | 37.5 |
| | | CB+ | 136.9 | 37.5 | 192.0 | 58.7 |
| | | Dual | 132.5 | 44.0 | 169.0 | 32.7 |
| | Ventrolateral | CR+ | 165.1 | 29.8 | 111.4 | 25.2 |
| | | CB+ | 148.8 | 35.1 | 133.0 | 27.1 |
| | | Dual | 126.9 | 45.9 | 122.3 | 39.1 |
| | Medial | CR+ | 134.2 | 27.8 | 128.6 | 35.7 |
| | | CB+ | 106.3 | 42.6 | 115.7 | 57.5 |
| | | Dual | 132.6 | 37.3 | 133.0 | 57.4 |
| Mid | Dorsolateral | CR+ | 132.4 | 27.0 | 164.7 | 48.8 |
| | | CB+ | 126.3 | 27.2 | 161.7 | 49.7 |
| | | Dual | 125.7 | 31.3 | 161.1 | 60.3 |
| | Ventrolateral | CR+ | 163.0 | 36.3 | 112.1 | 29.7 |
| | | CB+ | 172.0 | 33.9 | 124.8 | 30.0 |
| | | Dual | 142.6 | 28.4 | 116.5 | 34.1 |
| | Medial | CR+ | 143.6 | 31.8 | 116.7 | 27.7 |
| | | CB+ | 138.1 | 37.9 | 122.4 | 31.1 |
| | | Dual | 123.0 | 44.2 | 132.4 | 40.5 |
| Caudal | Dorsolateral | CR+ | 145.7 | 38.1 | 151.6 | 26.1 |
| | | CB+ | 114.5 | 50.4 | 131.6 | 42.0 |
| | | Dual | 126.8 | 39.1 | 136.3 | 37.7 |
| | Ventrolateral | CR+ | 134.6 | 38.2 | 135.4 | 31.8 |
| | | CB+ | 152.1 | 59.6 | 139.2 | 30.4 |
| | | Dual | 138.8 | 77.7 | 140.5 | 35.7 |
| | Medial | CR+ | 138.5 | 35.3 | 116.3 | 31.6 |
| | | CB+ | 128.4 | 42.0 | 101.3 | 41.5 |
| | | Dual | 100.9 | 28.7 | 87.7 | 43.0 |

*SD denotes standard deviation*

***Table S2. Mean cell area density of DAB CR$^{+}$ and CB$^{+}$ cells in all reuniens (RE) internal subregions across the rostro-caudal axis of the thalamus***

| **Axis** | **RE subregion** | **Calretinin ($CR^{+}$)** | | **Calbindin ($CB^{+}$)** | |
|---|---|---|---|---|---|
| | | **Mean** | **SD** | **Mean** | **SD** |
| Rostral | RE anterior (REa) | 7.8 | 3.0 | 9.2 | 1.6 |
| Rostromedial | RE median (REm) | 11.1 | 7.9 | 9.0 | 4.7 |
| | RE lateral (REl) | 6.7 | 3.6 | 22.0 | 18.4 |
| | RE dorsal REd) | 5.3 | 1.9 | 9.4 | 5.0 |
| | RE ventral (REv) | 7.8 | 1.2 | 16.3 | 9.9 |
| | RE other (REother) | 6.3 | 1.8 | 10.4 | 3.6 |
| Medial | RE caudal median (REcm) | 10.9 | 6.0 | 11.8 | 3.9 |
| | RE caudal dorsal (REcd) | 7.5 | 1.4 | 12.9 | 3.7 |
| | RE caudal posterior (REcp) | 6.2 | 1.1 | 8.6 | 1.9 |
| | Perireuniens (PRe) | 5.5 | 1.1 | 10.5 | 6.0 |
| Mediocaudal | RE caudal median (REcm) | 7.0 | 4.3 | 14.2 | 5.8 |
| | RE caudal dorsal (REcd) | 12.3 | 6.0 | 22.7 | 18.2 |
| | RE caudal posterior (REcp) | 16.0 | 17.7 | 14.3 | 13.8 |
| | Perireuniens (PRe) | 4.8 | 1.7 | 14.8 | 13.7 |
| Caudal | RE caudal posterior (REcp) | 8.8 | 5.2 | 14.4 | 10.9 |
| | RE caudal dorsal (REcd) | 12.5 | 7.4 | 22.2 | 14.6 |
| | Perireuniens (PRe) | 10.0 | 3.3 | 20.9 | 8.9 |

** means are in cells/.01mm$^2$*

*SD denotes standard deviation*
